# Maturation of striatal dopamine supports the development of habitual behavior through adolescence

**DOI:** 10.1101/2025.01.06.631527

**Authors:** Daniel J. Petrie, Ashley C. Parr, Valerie Sydnor, Amar Ojha, Will Foran, Brenden Tervo-Clemmens, Finnegan Calabro, Beatriz Luna

## Abstract

Developmental trajectories during the transition from adolescence to adulthood contribute to the establishment of stable, adult forms of operation. Understanding the neural mechanisms underlying this transition is crucial for identifying variability in normal development and the onset of psychiatric disorders, which typically emerge during this time. Habitual behaviors can serve as a model for understanding brain mechanisms underlying the stabilization of adult behavior, while also conferring risk for psychopathologies. Dopaminergic (DA) processes in the basal ganglia are thought to facilitate the formation of habits; however, developmental trajectories of habits and the brain systems supporting them have not been characterized *in vivo* in developing humans. The current study examined trajectories of habitual behavior from adolescence to adulthood and sought to understand how the maturing striatal DA system may act as a potential mechanism in the process of habit formation. We used data from two longitudinal studies (combined *n* = 217, 10 – 32 years of age, 1-3 visits each, 320 total sessions) to characterize normative developmental trajectories of basal ganglia tissue iron concentration (a proxy for DA-related neurophysiology) and goal-direct and habitual control behaviors in a two-stage decision-making task. Tissue iron concentrations across the basal ganglia and habitual responding during the two-stage sequential decision-making task both increased with age (all *p* < 0.001). Importantly, habitual responding was associated with tissue iron concentrations in the putamen (*F* = 4.34, *p* = 0.014), such that increases in habitual responding were supported by increases in putamen tissue iron concentration during childhood through late adolescence. Exploratory analyses of further subdivisions of anatomical regions found that this association was specific to the posterior putamen. These results provide novel evidence in humans that habitual behavior continues to mature into adulthood and may be supported by increased specialization of reward systems.

## Main

The transition from adolescence to adulthood involves developmental trajectories that establish stable, adult forms of operation (Larsen & Luna, 2018; Luna et al., 2015; Tervo-Clemmens et al., 2023). Understanding the brain mechanisms supporting this transition and the variability in normative brain-behavior development can clarify how deviations from these trajectories may contribute to psychopathology, which often begins to emerge during adolescence (Cicchetti, 2008; Cicchetti & Rogosch, 2002), and becomes more diagnosable in early adulthood (Collins & Muñoz-Solomando, 2018). Despite this, the brain processes that underlying behavioral stabilization during this period remain unclear. Whether normative or impaired, the stabilization of behavior is essential for the relative optimization of behavioral outcomes. Habit formation is an important part of normative development, reflecting the shift from variable to stable behavioral patterns from adolescence to adulthood. Thus, habit formation offers a model for investigating the brain mechanisms underlying the stabilization of developmental trajectories during this period. Adolescence also marks the maturation of mesocorticolimbic dopaminergic circuitry, including the basal ganglia (Wahlstrom et al., 2010), which is implicated in habit formation and habit-related psychopathologies such as obsessive-compulsive disorder (OCD) and substance use disorders (SUDs). Thus, the current study examined associations between normative developmental trajectories of habitual behavior and basal ganglia tissue iron, a marker of dopaminergic physiology that has been linked to cognitive development (Larsen, Bourque, et al., 2020; Larsen, Olafsson, et al., 2020; Larsen & Luna, 2015; Parr et al., 2022; Parr, Perica, et al., 2024).

Instrumental behavior can be classified as either goal-directed or habitual (Stimulus - Response, S-R; Balleine & Dickinson, 1998; Dickinson et al., 1995). Goal-driven behavior is engaged during action-outcome (A-O) learning, motivating behavior based on desired outcomes. When actions are repeated in the presence or the potential of strong reward, a habit can form. Once formed, habits can be defined as behaviors that persist even without the associated reward, making them outcome insensitive (Dickinson, 1985). This gradual process automates behavior by reducing reliance on goal-directed, executive resources, which then can inform the transition to adult stabilization including diagnosable psychopathologies. Although habits can be formed throughout life (Loewenstein et al., 2016), the ability to engage habitual systems continues to mature throughout adolescence into adulthood. For example, rodent studies indicate that adults can readily form habits, while adolescent remain predominantly goal-directed, evidenced by adjusting their actions based on the current value of outcomes (i.e., sensitive to outcome devaluation; Naneix et al., 2012; Rode et al., 2020; Serlin & Torregrossa, 2015; Simon et al., 2013; Towner et al., 2020). There is evidence for a similar pattern occurring in humans, including habitual alcohol use typically emerging in early adulthood, despite common experimentation during adolescence (Chan et al., 2008). Although executive control is available in adolescents (Ordaz et al., 2013; Simmonds et al., 2017), they engage it less consistently (Montez et al., 2017, 2019), leading to less efficient S-R learning and habit acquisition compared to adults.

Neural circuitry supporting goal-directed and habitual behaviors has been well delineated (Balleine & Dickinson, 1998; Balleine & O’Doherty, 2010; Yin et al., 2004, 2005; Yin & Knowlton, 2006); the transition from goal-directed to habitual behaviors is supported by a change from prefrontal and dorsomedial striatal (caudate in humans) function to motor-related cortical regions and dorsolateral striatal (putamen in humans) function, where dopaminergic afferents from the midbrain to the striatum are thought to partially mediate goal-directed and habitual control (Graybiel, 1990). Importantly, dopamine’s role in habit formation is underscored by evidence that disrupting dopaminergic neurons or their synaptic connections in the dorsolateral striatum impairs markers of habit acquisition and habit-related striatal activity (Faure et al., 2005; Hernandez et al., 2013; Wang et al., 2011). Conversely, increasing dopamine activity in the dorsolateral striatum has been shown to accelerate the development of habits, highlighting its key role in the transition to habit formation (Belin & Everitt, 2008; Ito et al., 2002; Malvaez & Wassum, 2018).

Studies directly investigating the role of dopamine in habit formation in developing healthy humans are limited by challenges in the use of positron emission tomography (PET) in pediatric populations. To address these limitations, our group has previously leveraged brain tissue iron as a non-invasive imaging marker of dopamine neurophysiology (Larsen, Olafsson, et al., 2020). Tissue iron is closely related to dopaminergic processes, particularly within the dopamine-rich basal ganglia, where high concentrations of brain iron are found (Brass et al., 2006; Connor et al., 1990; Hallgren & Sourander, 1958; Thomas et al., 1993). Tissue iron is involved in dopamine synthesis, and is a cofactor for tyrosine hydroxylase, a rate limiting enzyme in dopamine production (Ortega et al., 2007; Torres-Vega et al., 2012; Zucca et al., 2017). Iron has also been associated with regulating dopamine receptor function and neurotransmitter activity. Alterations of tissue iron have been linked to a wide range of disorders involving dopamine, including Parkinson’s (Moreau et al., 2018; Piao et al., 2017; Zucca et al., 2017), substance use disorders (Adisetiyo et al., 2019; Ersche et al., 2017; Raka et al., 2020; H. Tan et al., 2023), and Huntington’s disease (Ward et al., 2014), as well as neurodevelopmental disorders, such as Attention-Deficit/Hyperactivity Disorder (ADHD; Cascone et al., 2023).

Importantly, developmental increases in tissue iron accumulation have been observed in the basal ganglia over the first twenty years of life, after which the rate of accumulation decreases into adulthood (Aquino et al., 2009; Hallgren & Sourander, 1958; Hect et al., 2018; Larsen, Bourque, et al., 2020; Larsen, Olafsson, et al., 2020; Larsen & Luna, 2015; Peterson et al., 2019), and has been linked to cognitive performance (Larsen, Bourque, et al., 2020; Hect et al., 2018), functional connectivity in reward networks (Parr et al., 2021), and inhibitory control (Parr et al., 2022). Importantly, tissue iron can be measured non-invasively using magnetic resonance imaging (MRI), providing an indirect assessment of dopaminergic neurophysiology. Thus, increased tissue iron levels in regions like the striatum may serve as a marker for dopamine function, potentially reflecting changes related to aging, disease, or behavioral processes, such as habit formation.

In cognitive neuroscience, the distinction between habitual and goal-directed behaviors is often formalized as model-free and model-based control within reinforcement learning (RL) models, commonly assessed using the two-stage sequential decision-making task (Daw et al., 2005). In the task, participants make a sequence of two choices that lead to rewards, with the second-stage choice being probabilistically determined by the first-stage decision. Model-based control is computationally demanding, and involves forward planning where individuals use an internal model of the environment to predict outcomes based on the current state and update their decisions accordingly. This is thought to align with goal-directed behavior, as actions are performed with an understanding of their consequences. In contrast, model-free control relies on reinforcement from past rewards without considering future states, and is less computationally demanding, making it more reflexive and akin to habitual behavior. Although model-based and model-free control are often linked to goal-directed and habitual behaviors respectively, this relationship is not absolute, as both processes can influence decision-making simultaneously, and transitions between the two are dynamic rather than discrete (Lee et al., 2014).

The distinction between model-based and model-free control has been useful, but empirical evidence linking model-free control to habitual behavior is limited (Watson & de Wit, 2018). Most studies using the sequential decision-making task have found no association between model-free control and traditional habit indices (e.g., outcome devaluation; Friedel et al., 2014; Gillan et al., 2015; Sjoerds et al., 2016), and direct evidence supporting the link between model-free control and habits is lacking (Feher Da Silva & Hare, 2020). This has sparked debate on the independence of habits from model-free learning (Balleine et al., 2015; Balleine & Dezfouli, 2019; Ceceli & Tricomi, 2018; Daw et al., 2005; Dezfouli et al., 2014; Dezfouli & Balleine, 2012, 2013; Dolan & Dayan, 2013; Gershman et al., 2014; Miller et al., 2019; Morris & Cushman, 2019; Otto et al., 2013). Here, we leverage a specific stage of the two-stage task where we measure habits as repetitive behaviors that occur without conscious awareness of rewards, separating this construct from model-free learning given the latter’s reliance on reward contingencies. Thus, habitual behaviors in the current study reflect perseverative behavior regardless of task transitions or rewards. To this end, we consider a novel parameter from the two-stage decision making task—“first-stage stay”—to measure how often participants repeat the same first-stage choice across trials, irrespective of transition structures or previous rewards. As such, this form of perseveration serves as a measure of habit, as it represents rigid, automatic behavior that is insensitive to new information.

In order to better understand how maturation of the striatal dopamine system contributes to the maturation of these habitual behaviors, we used data from two longitudinal samples (*n* = 217, 10 – 32 years of age, 1-3 visits each, 320 total sessions), with up to three longitudinal visits per participant. We used generalized additive mixed models (GAMMs) to characterize non-linear normative developmental trajectories of tissue iron concentration across the basal ganglia, and developmental trajectories of model-based, model-free, and first-stage stay (habitual) behaviors. We then assessed whether normative trajectories of tissue iron concentration across the basal ganglia varied with individual differences in performance during the sequential decision-making task. We had two main hypotheses: 1) Based on rodent models of habitual control (Naneix et al., 2012; Rode et al., 2020; Serlin & Torregrossa, 2015; Simon et al., 2013; Towner et al., 2020), habitual behaviors (operationalized as first-stage stays) will increase in adolescence and stabilize into adulthood. 2) Given that the transition from goal-directed to habitual behavior is supported by increased reliance on the putamen (Amaya & Smith, 2018; Foerde, 2018; Knowlton & Patterson, 2018; Patterson & Knowlton, 2018; Tricomi et al., 2009), we hypothesized that developmental increases in putamen tissue iron, as a putative marker of DAergic neurophysiology, would support age-related increases in first-stage stays, with adults and adolescents who expressed a higher degree of habitual behavior showing higher levels of putamen tissue iron relative to their same-aged peers. In support of these hypotheses, this study provides initial evidence that the maturation of dopaminergic input to the posterior putamen supports the emergence of habitual behavior during the adolescent period and may reflect the stabilization of cognitive processes.

## Results

### Zero-order correlations and descriptive statistics

Zero-order correlations between developmental, behavioral, and neurobiological features are presented along with descriptive statistics in Table 3. All behavioral measures were positively associated with each other (*r*’s 0.60 – 0.71), indicating that model-based, model-free, and first-stay stay terms were linked. All tissue iron measures were positively associated (*r*’s 0.06 – 0.65), indicating tissue iron coupling in the basal ganglia across persons. Age was significantly correlated with all behavioral and tissue iron measures except the caudate (|*r*| 0.04 – 0.56) as seen previously (Parr et al., 2022).

### Striatal tissue iron concentrations increase with age

Age-related differences in nT2*w for all regions were significant at a Bonferroni-adjusted significance threshold of α = 0.0125, controlling for sex and visit number (Table S1). nT2*w regional distributions (Figure 3A) and developmental trajectories (Figure 3B) are similar to those reported in prior studies (Larsen, Olafsson, et al., 2020; Parr et al., 2022; Peterson et al., 2019; Price et al., 2021). Tissue iron concentration steadily increased with age in the globus pallidus (*edf* = 1.21, *F* = 43.68, *p* < 0.001), nucleus accumbens (*edf* = 1, *F* = 18.69, *p* < 0.001), and putamen (*edf* = 1, *F* = 87.16, *p* < 0.001). We did not find monotonic tissue iron increases in the caudate, although the age smooth was significantly different than 0 (*edf* = 1.92, *F* = 5.94, *p* = 0.003). Tissue iron is inversely related to nT2*w, such that smaller values reflect more tissue iron and larger values reflect less tissue iron. For ease of visual interpretation, figure axes for tissue iron have been reversed such that high tissue iron (low nT2*w) is higher on figure axes. Measures of tissue iron for each ROI were included in subsequent analyses as an index of striatal dopamine neurobiology.

**Figure 1.**
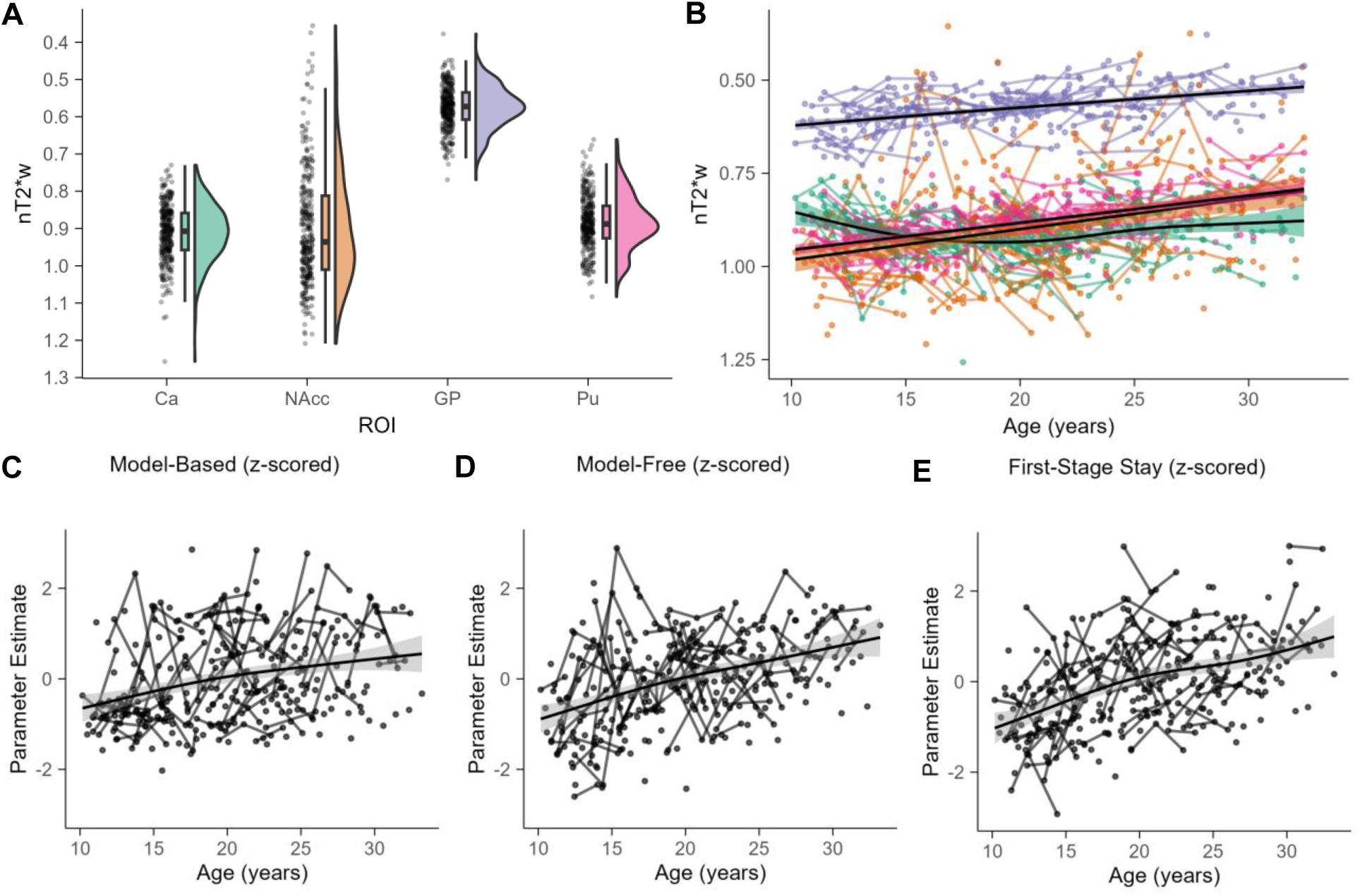
Developmental trajectories for tissue iron and behavioral measures. Distributions (A) and developmental trajectories (B) of nT2*w in basal ganglia structures. These findings replicate previous developmental work. In A and B, the y-axis is reversed such that lower values reflect more tissue iron. All age smooths are significantly different (all *p* < Bonferroni-adjusted α = 0.0125) than 0 and increased over age. (C) The model-based effect is plotted as a smooth function and reflects the reward by transition interaction fixed effect plus the reward by transition random interaction random effect from a multilevel logistic regression model with age excluded. (D) The model-free effect is plotted as a smooth function and reflects the main effect of reward as a fixed effect plus the main effect of reward as a random effect. (E) The first stage stay effect is plotted as a smooth function and reflects the intercept plus the random intercept. The gray band represents 95% CI.

**Figure 2.**
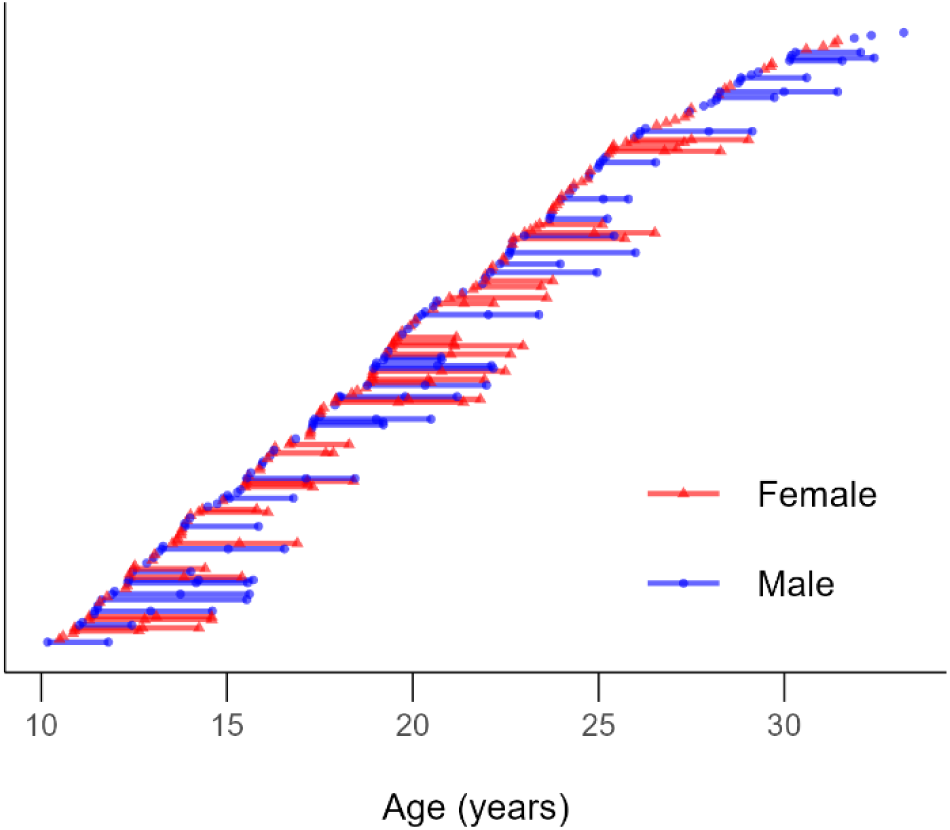
Age range of final sample. 320 scans were nested in 217 individuals. Each point depicts an individual session, with lines connecting individual participants across sessions.

**Figure 3.**
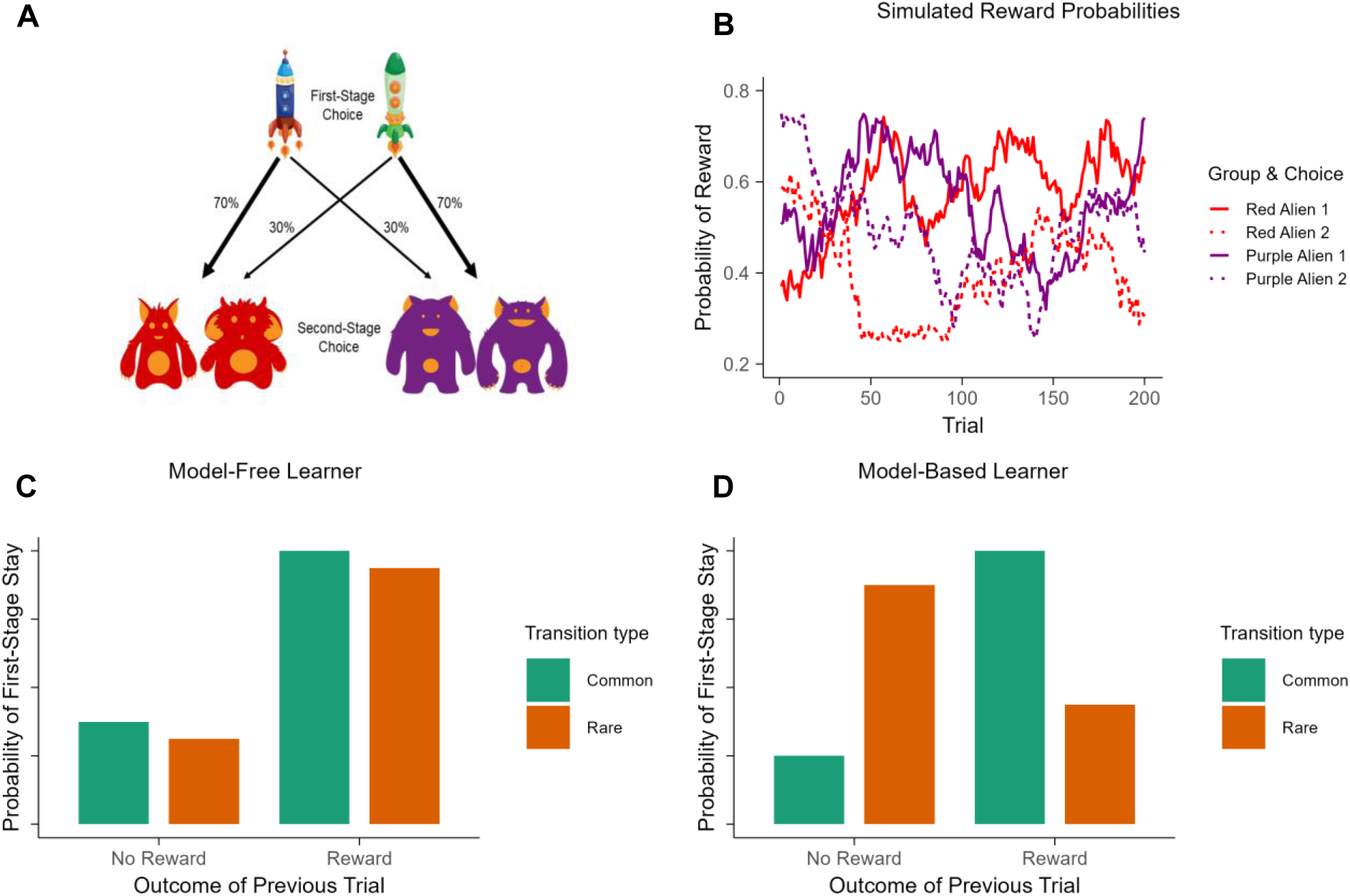
Design of the two-stage sequential decision-making task and hypothetical model-free and model-based behavior. (A) On each trial, participants first choose between two spaceships (first-stage choice) leading to a probabilistic transition (either 70% or 30%) to either a red planet or a purple planet. Then, participants select between two aliens (second-stage choice) and are either rewarded with space treasure or not. (B) The probability of winning space treasure is shown as a function of trials for each alien during the second-stage choice. Reward probabilities for the four choices changed independently via Gaussian random walks (*mean* = 0, *SD* = 0.025) with boundaries set to 0.25 and 0.75. (C) Model-free learning theory predicts that a first-stage choice that results in a reward is more likely to be repeated on the next trial, regardless of transition type. (D) Model-based learning theory predicts that a rare transition should influence the perceived value of the other first-stage option, leading to an interaction effect between reward and transition type.

### Measures of two-stage sequential decision-making tasks increase with age

All behavioral age smooths were significant at a Bonferroni-adjusted significance threshold of α = 0.0167, controlling for sex and visit number. Results from the analyses examining age related change in measures from the two-stage sequential decision-making task are presented in Figure 3C—3E and Supplementary Table S2. In line with past literature (Decker et al., 2016; Vaghi et al., 2020), we observed a significant increase of model-based behavior with age (Figure 4A.; *edf* = 1.49, *F* = 15.29, *p* < 0.001), indicating that participants tended to increasingly integrate knowledge of the task structure with age. We also observed a significant increase in model-free behavior with age (Figure 4B.; *edf* = 1.13, *F* = 52.45, *p* < 0.001), indicating that the probability of repeating a first-stage stay after a reward increased with age. First-stage stay behavior—our measure of habit—also increased with age (Figure 4C.; *edf* = 1.71, *F* = 40.13, *p* < 0.001), indicating that older participants were more likely to repeat the same choice at the start of each trial regardless of transition-type or whether or not they were previously rewarded.

**Figure 4.**
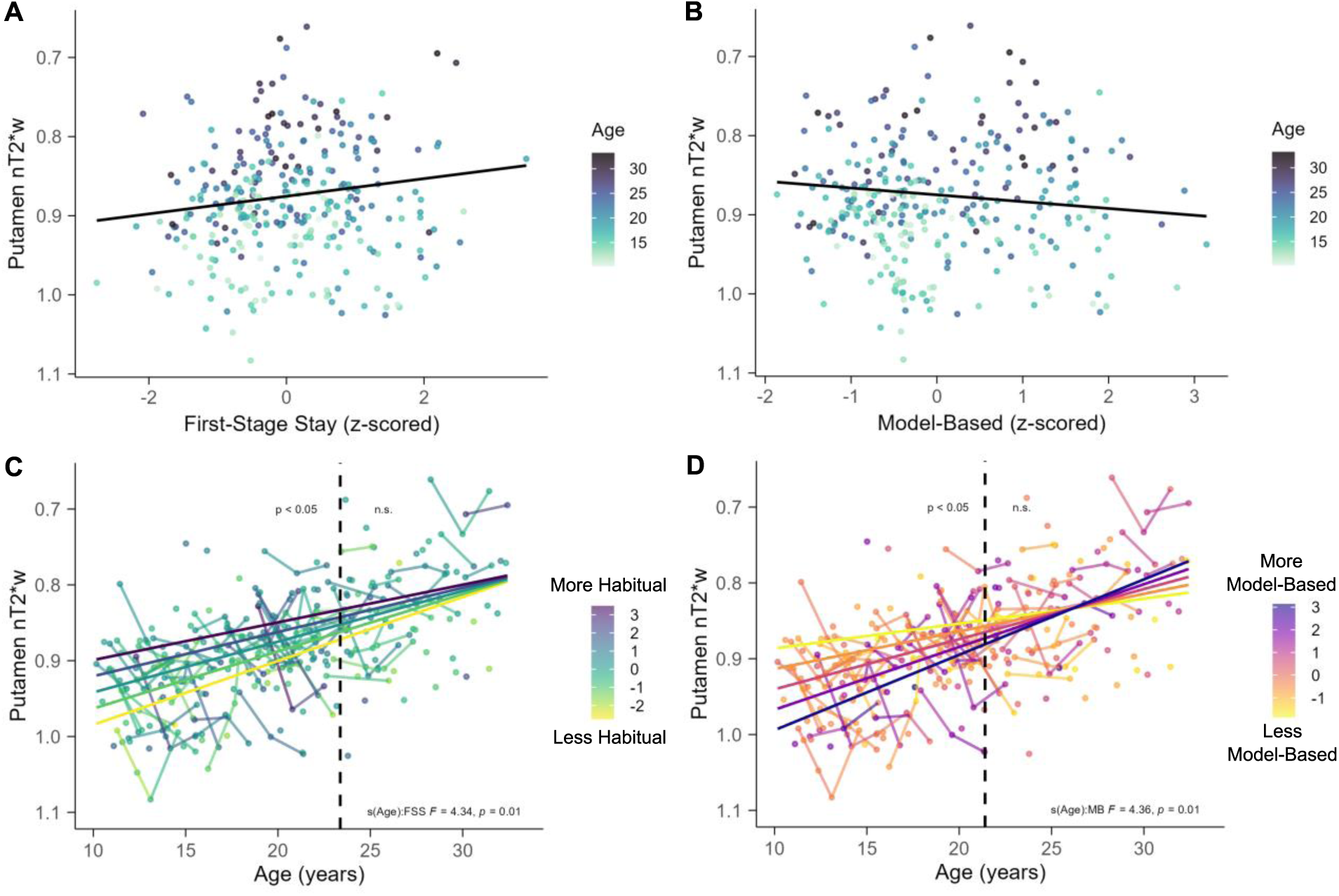
GAMM results for Model 3 and time-varying parameter results with putamen tissue iron. In all panels, the y-axis is reversed such that lower values reflect more tissue iron (A) There was a significant positive association between first-stage stay behavior and putamen tissue iron, such that more first-stage stays were associated with higher levels of putamen tissue iron. (B) There was a significant negative association between model-based behavior and putamen tissue iron, such that more model-based behavior was associated with lower levels of putamen tissue iron. In (A) and (B) the color of the points represents age of the participant with darker colors corresponding with older ages. (C) First-stage stay behavior moderated the association between age and putamen tissue iron. The dashed vertical line at 23.37 years old reflects the age boundary where the 95% CI of the first derivate does not include 0, indicating significance. (D) Model-based behavior moderated the association between age and putamen tissue iron. The dashed vertical line at 21.40 years old reflects the age boundary where the 95% CI of the derivate does not include 0, indicating significance.

We conducted three supplementary analyses to maintain consistency with developmental studies using the task, facilitating comparability across future research (Decker et al., 2016; Nussenbaum et al., 2020; Potter et al., 2017). First, we fitted a multilevel logistic regression model where age was included as a main effect along with all possible two- and three-way interactions between task variables. Results generally match other research groups (Decker et al., 2016; Nussenbaum et al., 2020; Potter et al., 2017). We found that across age, participants used a combination of behavioral strategies (i.e., the main effect of reward and the interaction between reward and transition type were both significant), and that adults appear to be using all strategies at higher level than adolescents. Second, we also assessed whether participant responses reflected knowledge of task structure. Results from this multilevel model suggested that participants were slower to respond during the second-stage choice after rare transitions compared to common ones (i.e., a main effect of transition type), suggesting that participants were overall aware of the task structure. The difference in response times after rare and common transitions increased across the age range. Third, we assessed whether knowledge of task structure was related to model-based control. Results from this multilevel model suggest that slower reaction times after rare transitions were associated with more model-based control, however, early adolescents do not appear to be integrating this information to the extent the mid/late adolescents and adults do. We present the full results from these analyses in the supplementary materials (section S.6.1 and section S.6.2).

### Habitual responding is associated with elevated putamen tissue iron

Because all behavioral and tissue iron measures increased with age, which could lead to concurvity (a form of multicollinearity) issues when model fitting within a GAM framework, we chose to regress out age-related effects from the behavioral parameters to orthogonalize the age smooth and the smooths associated with behavioral performance. Age-residualized estimates of the behavioral parameters were obtained by linearly regressing age onto each behavioral parameter. These age-residualized estimates of behavioral performance were used for all subsequent models.

To determine the best fitting model for identifying behavioral measures that parsimoniously capture variability in tissue iron above and beyond age, we compared a set of four models (see Table 2) for each ROI using analysis of deviance tests and AIC comparisons. If the results from both the analysis of deviance tests and the AIC index favored a more complex model, we retained it for further examination. The full results from these analyses are presented in Supplementary Tables S3 – S7. For the putamen ROI, both the AIC and the analysis of deviance tests indicated that Model 3, which included smooths for first-stage stay and model-based behavior, was the best fitting model, supporting associations between putamen tissue iron and both of these behaviors. For all other ROIs, Model 1, which included only an age effect and no behavioral measures, was determined to fit the data best, suggesting that performance during the task did not provide significant improvement in explaining variability in tissue iron trajectories.

Next, we examined the main effects specified in Model 3 for the putamen. Results indicated that increased first-stage stay behavior was associated with increased putamen tissue iron (Figure 4A.; *edf* = 1, *F* = 6.21, *p* = 0.01) controlling for age. Interestingly, the association between model-based learning and putamen tissue iron showed the opposite effect, such that increased model-based learning was associated with decreased putamen tissue iron (Figure 4B.; *edf* = 1, *F* = 4.14, *p* = 0.04) controlling for age.

### Habitual behavior moderates developmental trajectories of putamen tissue iron

Next, we investigated whether developmental trajectories of tissue iron varied with behavioral performance, with the goal of understanding whether the maturation of dopamine-related neurophysiology diverges between individuals who express different degrees of habitual behavior. We found that first-stage stay (*edf* = 2, *F* = 4.34, *p* = 0.01) and model-based behavior (*edf* = 2, *F* = 4.36, *p* = 0.01) significantly moderated developmental trajectories of putamen tissue iron in interaction analyses (see Table S8).

For first-stage stay behavior, analysis of the derivative of the fitted age trajectory indicated that there was a significant interaction between age and first-stage stay behavior on putamen tissue iron between 10.17 and 23.37 years old (dashed line, Figure 4C). Marginal effects between age and tissue iron were extracted at representative fixed values of ±2 SD from the mean, ±1 SD from the mean, and the mean amount of first-stage stay behavior, and plotted for visualization purposes (Figure 4C). Individuals who had more first-stage stays for their age had elevated putamen tissue iron compared to individuals who had fewer first-stage stays for their age. This effect was more pronounced at younger ages and decreased in magnitude across age within the significant age range.

For model-based behavior, analysis of the derivative of the fitted age trajectory indicated that the interaction between age and model-based behavior on putamen tissue iron was significant between 10.17 and 21.40 years old (dashed line, Figure 4D). Similar to first-stage stay behavior, marginal effects between age and tissue iron were extracted at fixed values of 0, ±1, and ±2 SD from the mean amount of model-based behavior, and plotted for visualization purposes (Figure 4D). Individuals with higher values of model-based behavior for their age had lower putamen tissue iron compared to individuals who had lower values of model-based behavior for their age. This effect was also more pronounced at younger ages and decreased in magnitude across age within the significant age range. Taken together, results from the time-varying effects models suggest that, for younger participants, individuals with heightened putamen tissue iron respond more habitually and are less model-based relative to low putamen tissue iron.

### Exploratory analyses focusing on subdivisions of putamen

Projections from the cortex to the putamen are topographically organized such that prefrontal areas project to more anterior parts of the putamen, and sensorimotor and motor areas project to more posterior parts of the putamen (Haber, 2016). Thus, different areas of putamen have been implicated in different functions, with more anterior regions being previously associated with goal-directed responding during instrumental learning tasks (Akkermans et al., 2018; Sjoerds et al., 2013), and more posterior regions being previously associated with establishing more habitual stimulus-response associations (de Wit et al., 2012; Gillan, Robbins, et al., 2016; Tricomi et al., 2009). Considering these functional differences, we were interested in examining which subdivisions of the putamen were driving the significant associations between age, behavior, and tissue iron. We hypothesized that posterior putamen would be driving the positive association between first-stage stays and tissue iron, due to the outcome insensitive (i.e., non-goal directed) aspects of the first-stage stay measure, and its potential role in the engagement of automatic motor responses in habitual processes.

To test the hypothesis, we extracted tissue iron data from an anatomically-defined atlas which has been previously validated based on dopaminergic Positron Emission Tomography (PET; [11C]PHNO) data (Tziortzi et al., 2011), and which has been used to more finely parcellate striatal subdivisions, including in a previous study from our group (Calabro et al., 2023). This atlas separates putamen into four parts: anterior dorsal putamen, anterior ventral putamen, posterior dorsal putamen, and posterior ventral putamen. Distributions of tissue iron and their developmental trajectories across the four parts of putamen were similar to the full-putamen results and did not show any regional differences (Figure 5; all age smooths *p* < 0.001). We then fit a series of models similar to Model 3 (see Table 2 and Equation 3), to assess which areas of the putamen were significantly associated with age, first-stage stays, and model-based learning. We applied the Bonferroni correction across these sets of models to account for multiple comparisons across the four subsections of bilateral putamen, resulting in a significance threshold of α = 0.0125.

**Figure 5.**
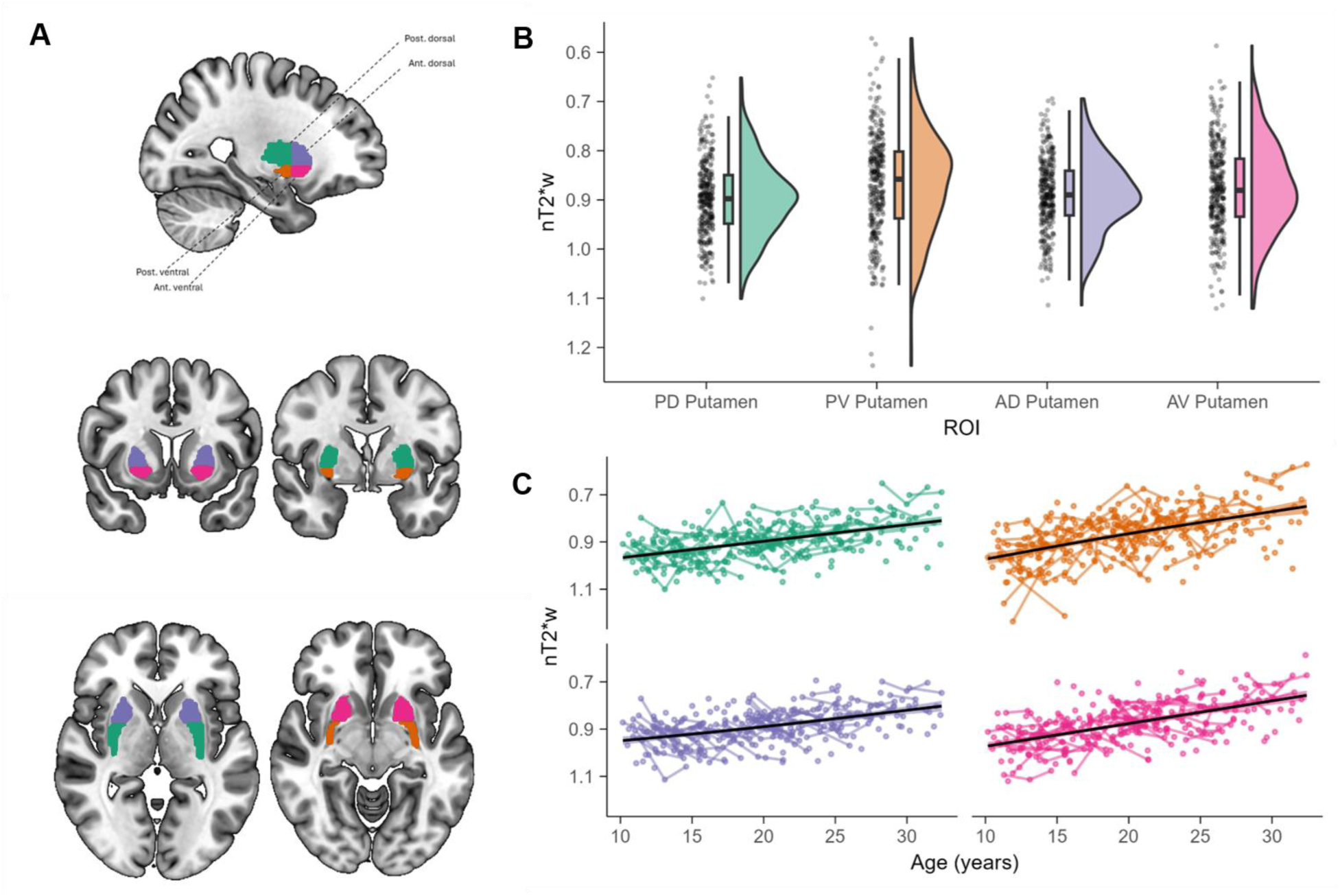
(A) Subdivisions of the putamen used for exploratory analyses. Distributions (B) and developmental trajectories (C) of putamen nT2*w. In B and C, the y-axis is reversed such that lower values reflect more tissue iron. All age smooth functions are significantly different than 0 and increased over age.

Results from exploratory time-varying parameters analyses are presented in Figure 6. For anterior regions of the putamen, there was no evidence of a time varying effect for first-stage stay behavior (Figure 6A & 6C; *p*-values > Bonferroni-adjusted α = 0.0125). However, for both posterior regions of putamen, first-stage stay behavior moderated the association between age and tissue iron (Figure 6B posterior dorsal putamen: *edf* = 2.63, *F* = 3.37, *p* = 0.02; Figure 6D posterior ventral putamen: *edf* = 2.79, *F* = 3.84, *p* = 0.0124), although only the effects in the posterior ventral putamen reached the Bonferroni-adjusted α of 0.0125.

**Figure 6.**
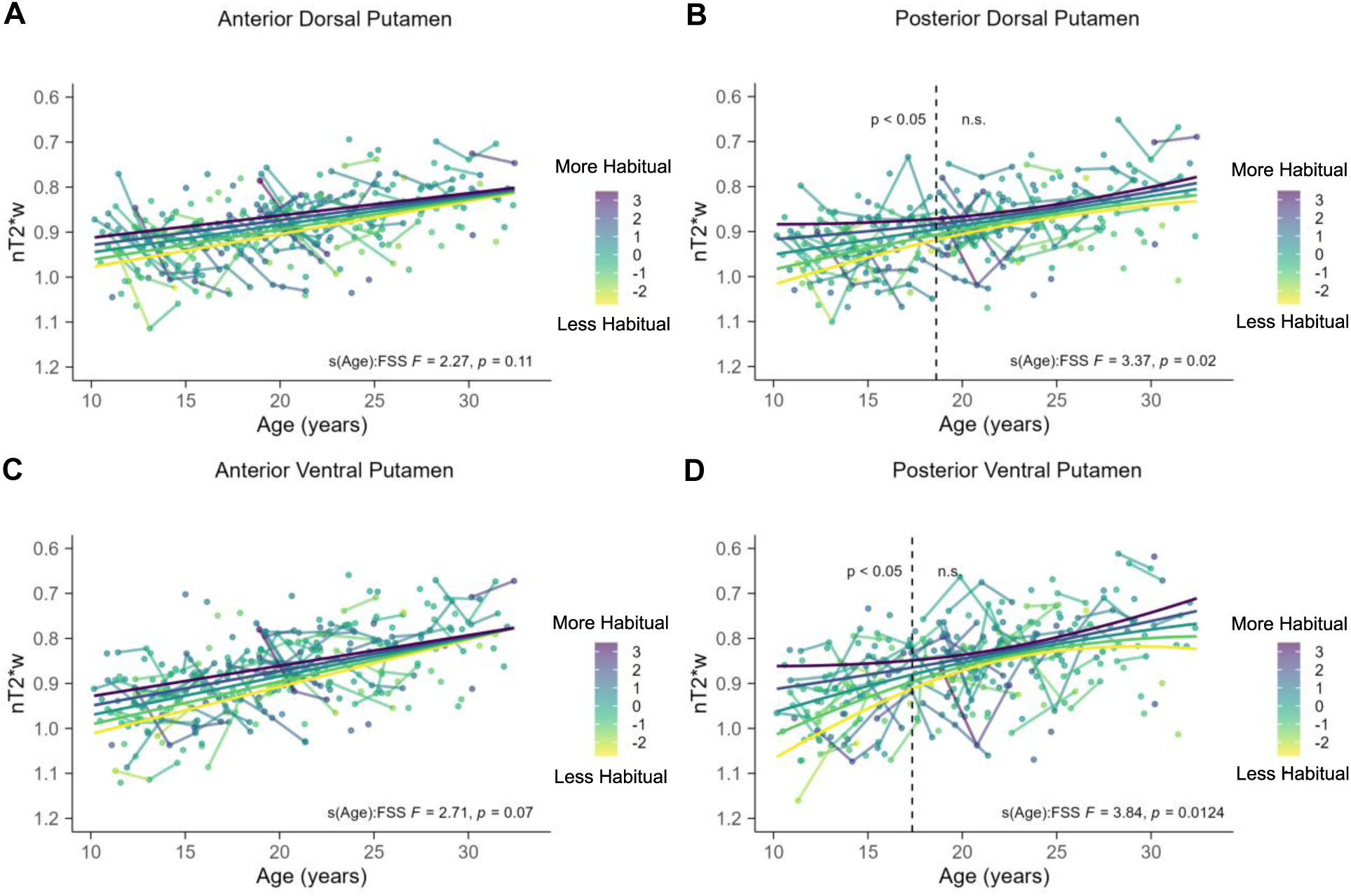
Time-varying parameter model results focusing on first-stage stay. In all panels, the y-axis is reversed such that lower values reflects more tissue iron. The color of the points represents first-stage stay z-scores with darker colors indicating more first-stage stays. There were no significant associations between age and first-stage stay behavior on anterior dorsal putamen tissue iron (A) or anterior ventral putamen tissue iron (C). (B) First-stage stay behavior moderated the association between age and posterior dorsal putamen tissue iron. The dashed vertical line at 18.62 years old reflects the age boundary where the 95% CI of the derivate does not include 0, indicating significance. (D.) First-stage stay behavior moderated the association between age and posterior ventral putamen tissue iron. The dashed vertical line at 17.35 years old reflects the age boundary where the 95% CI of the derivate does not include 0, indicating significance.

For posterior dorsal putamen, analysis of the derivative of the fitted age trajectory indicated that the interaction between age and first-stage stay behavior on tissue iron was significant between 10.17 and 18.62 years old (dashed vertical line Figure 6B). For posterior ventral putamen, analysis of the derivative of the fitted age trajectory indicated that the interaction between age and first-stage stay behavior on tissue iron was significant between 10.17 and 17.35 years old (dashed vertical line Figure 6D). Marginal effects between age and tissue iron were extracted at fixed values of ± 2 *sd* from the mean, ± 1 *sd* from the mean, and the mean amount of first-stage stay behavior, and plotted for visualization purposes.

Taken together, results from the exploratory analyses suggest that the time-varying effects of first-stage stay behavior may be more localized to the posterior putamen, and are similar to the time-varying effects observed in the main analyses for the putamen as a whole. Adolescents who had more first-stage stays for their age had elevated posterior putamen tissue iron compared to individuals who had less first-stage stays for their age. This effect was more pronounced at younger ages, and decreased in magnitude across age within the significant age range. See Table S9 for the table with full results and Figure S1 for the model-based results.

## Discussion

In this study, we examined the maturation of basal ganglia tissue iron, performance during a sequential decision-making task, and their interaction, in a sample of 217 adolescent and young adult participants assessed across 1-3 longitudinal timepoints. We leveraged a novel measure of perseverative responses reflecting habit (first-stage stay) that has not been considered in the developmental literature, to demonstrate increased reliance on habitual decision making from adolescence to adulthood. We also replicated previous work characterizing the normative developmental trajectories of tissue iron in the basal ganglia, and work suggesting that model-based behaviors increased with age. Developmental increases were found ubiquitously across behavioral strategies, underscoring adult abilities to transition between strategies more efficiently than adolescents. Finally, we present novel evidence that variation in developmental trajectories of putamen tissue iron, particularly in more posterior regions, were associated with individual differences in habitual and model-based responding, such that individuals with higher iron in the putamen, reflecting maturation of DAergic neurophysiology, responded more habitually and less goal-directed during late childhood and through adolescence.

Similar to other developmental studies that have administered the sequential decision-making task, we found that model-based control increased with age (Decker et al., 2016; Nussenbaum et al., 2020; Potter et al., 2017; Scholz et al., 2023; Vaghi et al., 2020). We also observed increases in the tendency to repeat a first-stage choice with age, consistent with other research findings (Decker et al., 2016; Potter et al., 2017). Importantly, we add to this literature by computationally validating first-stage choices as a developmentally sensitive marker of habitual behavior in a two-stage goal-directed task. This perseverative behavior was on average low in children but increased with age, aligning with less response variability seen in older participants (Christakou et al., 2013) and with findings that adolescents are less consistent in engaging specific brain processes (Montez et al., 2017, 2019). Although ample evidence suggests that adult-like behavior is present during adolescence (Ordaz et al., 2013; Simmonds et al., 2017), the greater response variability observed in childhood and early adolescence may reflect an adaptive and exploratory search process (Parr, Sydnor et al., 2024). This developmental bias likely promotes environmental exploration and optimal responding as individuals mature into adulthood (Gopnik et al., 2015). Across studies, developmental increases in perseverative behavior may index the transition from more exploratory, flexible responses to more rigid and efficient ones (Parr, Sydnor et al., 2024) such as a readiness to generate habitual responses. Rodent models have shown that adolescents have difficulty forming habits compared to adults (Naneix et al., 2012; Rode et al., 2020; Serlin & Torregrossa, 2015; Simon et al., 2013; Towner et al., 2020); here we demonstrate a similar developmental profile, for the first time in humans.

We did not find any association between putamen tissue iron measures and the model-free parameter, which is generally in line with the literature using the sequential decision-making task in adults. Most studies have found no association between model-free control and traditional indices of habit (e.g., outcome devaluation; Friedel et al., 2014; Gillan et al., 2015; Sjoerds et al., 2016), dopaminergic alleles in the striatum (dopamine- and cAMP-regulated neuronal phosphoprotein, DARPP-32) and prefrontal cortex (Catechol-O-methyltransferase, COMT; Doll et al., 2016), cortisol response to acute stressors (Otto, Raio, et al., 2013), substance use disorders (Hogarth, 2020; Sebold et al., 2014), and placebo versus drug enhancing dopamine treatment (L-DOPA; Wunderlich et al., 2012). Importantly, and in contrast to habits, rats and humans show decreases in model-free control as a result of extensive training on the sequential decision-making task (Economides et al., 2015; Miller et al., 2017). These findings suggest that model-free control may be limited in fully capturing biological or psychological constructs that are usually associated with habit. Alternatively, model-free responses may reflect *habit formation*, as it emphasizes the development of reward-based, automated responses that are sensitive to reward contingencies during initial learning. In contrast, the first-stage stay parameter may be reflecting *habit perseverance*, or the persistence of rigid, automatic S-R behavior that is insensitive to new information (e.g., outcome values or transition probabilities). Habits can be defined as repetitive behaviors that are outcome-insensitive (de Wit et al., 2018). Thus, perseveration, operationalized here as the tendency to stay at the same first-stage choice across trials, could serve as an additional measure of habit. Unlike model-free control, which relies on conscious reward-driven learning, perseverative behavior reflects habitual responding that occurs irrespective of reward or transition type during the task. This distinction highlights that model-free control may index the formation of habitual patterns, while first-stage stay behavior captures the persistence of these patterns over time.

An important question, and avenue for future study, is how the brain determines when to rely on each learning strategy and whether the arbitration between these strategies differs developmentally. Some research groups suggest that humans use meta-decision-making (the process of selecting between different learning strategies) when deciding between strategies (Boureau et al., 2015; Gershman et al., 2015; Kool et al., 2016). In a stable environment, a meta-rational agent would likely use the least effortful approach, favoring a perseverative or more automatic strategy like a first-stage stay. As the environment becomes more complex, a meta-rational agent would weigh the costs and benefits of exerting the extra effort required for more complex strategies (e.g., model-free, or model-based). In reality, humans employ a wide variety of learning strategies (Feher Da Silva & Hare, 2020; Momennejad et al., 2017), and future work should continue to characterize these strategies’ relative contributions and developmental patterns. Our findings that first-stage stay, model-based, and model-free parameters all increased with age may be supported by known improvements in executive function into adulthood (Tervo-Clemmens et al., 2023) allowing the ready ability to make stimulus response association facilitating the transition to habitual responding as well as allowing for more efficient switching between effortful and automatic strategies compared to adolescents or children.

Our findings of increases in tissue iron in the striatum align with past research indicating that striatal tissue iron levels increase throughout adolescence, and play a role in cognitive development including inhibitory control and risk taking, as well as specialization of frontostriatal reward-related connectivity (Larsen, Bourque, et al., 2020; Parr et al., 2021, 2022). Adding to this literature, we found that tissue iron in the putamen, a region central to habit formation (Foerde, 2018), is associated with individual differences in habitual behavior. Specifically, increased first-stage stays were associated with high putamen tissue iron levels through late adolescence and into early adulthood. These results support the hypothesis that dopaminergic processes, indexed by tissue iron, play a role in stabilizing behavior and facilitating habit formation from adolescence into adulthood. Dopaminergic signaling in the striatum, particularly the putamen, is known to strengthen S-R associations, promoting neuroplasticity and specialization within corticostriatal circuitry (Horvitz, 2009). During adolescence, enhanced dopamine function in the striatum may contribute to more reward-driven behavior and resistance to habit formation (Larsen & Luna, 2018; Luciana et al., 2012; Spear, 2000). Developmental increases in dopamine-related processes contribute to the specialization of corticostraital circuitry into adulthood (Parr et al., 2021), which may play a role in the encoding of well-learned S-R associations that are critical for the automaticity of habitual actions. In the current study, adolescents who demonstrated more habitual responses had heightened tissue iron compared to those who exhibited less habitual behavior, potentially indicating more advanced maturation of striatal circuitry or reflecting trait-level differences in dopaminergic function that may facilitate increased habitual behavior.

Our exploratory analyses support the role of the posterior putamen in habitual processes, consistent with the organization of cortical projections to the putamen. Specifically, anterior regions of the putamen receive input from associative areas of the frontal cortex, while posterior regions receive input from the primary motor cortex and the supplemental motor area (Alexander et al., 1986; Haber, 2003; Parent & Hazrati, 1995). Thus, different areas of putamen support distinct functions, with anterior regions associated with goal-directed responding during instrumental learning tasks (Akkermans et al., 2018; Sjoerds et al., 2013), and posterior regions previously linked to habitual stimulus-response associations (de Wit et al., 2012; Gillan, Robbins, et al., 2016; Tricomi et al., 2009). Our results align with these functional distinctions; we found that associations between tissue iron and habitual responding were specific to the posterior putamen during adolescence. This functional differentiation suggests that as the adolescent brain develops, dopaminergic systems in the posterior putamen may bias certain behaviors toward habitual responding, which could have implications for understanding how brain organization supports the transition from goal-directed to habitual control during this critical developmental period.

Findings from this study may inform the development of compulsive psychopathologies, such as obsessive-compulsive disorder (OCD) and substance use disorders (SUDs), both characterized by rigid, maladaptive behaviors that persist despite negative outcomes (Everitt & Robbins, 2005). Compulsions and habits share neurobiological and psychological features, where many compulsive disorders are initially driven by goal-directed actions that shift to outcome-independent, stimulus-response (S-R) habits (Dayan, 2009; Everitt & Robbins, 2005). Research indicates that individuals with OCD often exhibit deficits in goal-directed control and heightened habitual control compared to non-clinical participants (Gillan et al., 2011, 2014; Gillan, Kosinski, et al., 2016; Voon, Baek, et al., 2015; Voon, Derbyshire, et al., 2015). Habit-related corticostriatal connectivity has also been linked to increased obsessive-compulsive symptoms during adolescence (Petrie et al., 2024), and may signal risk for OCD onset. Similarly, in SUD models, habitual and rigid behaviors are supported by dorsal striatal activity in drug exposed rodents (Corbit et al., 2012; Dickinson et al., 2002; Nelson & Killcross, 2006; Schmitzer-Torbert et al., 2015; Zapata et al., 2010). In human studies, heavy alcohol use has been associated with impairments in model-based control and increases in habitual control, which may contribute to the maintenance of addiction (Voon et al., 2017). These findings add to the existing literature and offers insight into the neurodevelopmental mechanisms underlying compulsive psychopathologies. This adolescent shift toward habitual control may predispose individuals to compulsive behaviors, where an early reliance on outcome-independent habits may solidify maladaptive behavioral patterns that are difficult to modify.

The current study has several limitations. First, we did not ask participants about their knowledge of the task structure. Therefore, we are unable to ascertain whether adolescents formed a cognitive model of the task (i.e., the ability to distinguish common versus rare transitions), and whether or not they used this knowledge to influence first-stage choices. However, we assessed reaction times following rare versus common transitions to gauge participants’ understanding of task structure, with slower responses expected after rare transitions if they understood the task (Decker et al., 2016; Nussenbaum et al., 2020; Potter et al., 2017). As shown in Supplementary Section 6.1.2, participants were indeed slower after rare transitions, indicating awareness of the task structure. Second, the first-stage stay parameter has not yet been validated against other habit tasks, habit questionnaires, or real-life habitual behavior, although we did demonstrate that it correlates with the RL model perseveration parameter, suggesting that it is tapping into related processes. Future studies should examine bivariate relations to other paradigms that are though to index habits (e.g., outcome devalue, slips-of-action, Pavlovian-to-Instrumental Transfer). Third, the use of nT2*w as a measure of dopaminergic processes was limited to the basal ganglia, given that this region is high in iron and we can therefore attribute signal to iron content there relative to other cortical areas where myelin and other critical neurodevelopmental processes may contribute to the signal. Therefore, this precludes our examination of other brain regions that receive dopaminergic projections and may be involved in habit formation.

In conclusion, this study provides novel evidence in humans that the ability to form habitual responses continues to mature into adulthood and may be underlied by the well-established maturation of striatal dopaminergic function. Specifically, we found that as tissue iron increases with age in the putamen, a key region for supporting habits, habit formation was enhanced and individual differences in tissue iron corresponded with more perseverative (habitual) behavior and less model-based behaviors. Understanding how changes in tissue iron concentrations contribute to the formation of habits and habitual behavior from adolescence to adulthood may provide insight into the emergence of psychopathologies related to compulsive habits that begin to emerge during this time like OCD and SUDs as well as psychopathologies more broadly as they all engage maladaptive habitual processes.

## Methods

### Participants

Participants ranging from 10 – 33 years old were drawn from two longitudinal developmental neuroimaging studies from our group (“Study A” and “Study B”) with data harmonized across studies for analyses (*n* = 217, 53% female, 1-3 visits, total visits = 320 across both studies, Figure 1). Both studies used an accelerated longitudinal design, recruiting participants with a uniform age distribution, balanced for biological sex assigned at birth, with up to three visits per participant (approximately 18 months apart). Study A included behavioral data from a two-stage sequential decision-making task (see Task Methods) and fMRI data at all three visits. Therefore, all available data from Study A were used (*n* = 156, age-range = 10.17 – 32.42, 51% female, 1 – 3 visits, total visits = 259). Study B collected fMRI and two-stage sequential decision-making task data at one time point (i.e., cross-sectional; *n* = 61, age-range = 13.65 – 33.22, 56% female, 1 visit, total visits = 61). Participants from both samples were recruited from the community and screened for the following criteria: no recent loss of consciousness, no self or first-degree relatives with major psychiatric or neurological diagnoses, an IQ score below 80, and no MRI contraindications (e.g., non-removable metal in the body, claustrophobia).

Demographic information for the current study can be found in Table 1. The University of Pittsburgh’s Institutional Review Board approved both studies and both complied with the Code of Ethics of the World Medicine Association (Declaration of Helsinki, 1964). Adult participants (>18) provided informed consent. For minors, parents provided consent and youths under 18 provided assent. Participants were compensated for completing the assessments.

**Table 1.** Zero-order correlations and descriptive statistics of all variables in the current study.

| Variable | 1 | 2 | 3 | 4 | 5 | 6 | 7 | *M* | *SD* |
| --- | --- | --- | --- | --- | --- | --- | --- | --- | --- |
| 1. Age |  |  |  |  |  |  |  | 20.23 | 5.64 |
| 2. Model-based | 0.33* |  |  |  |  |  |  | 0.40 | 0.39 |
| 3. Model-free | 0.45* | 0.60* |  |  |  |  |  | 0.37 | 0.26 |
| 4. First-stage stay | 0.48* | 0.71* | 0.71* |  |  |  |  | 1.46 | 1.06 |
| 5. Globus pallidus tissue iron | -0.45* | -0.16* | -0.17* | -0.24* |  |  |  | 0.57 | 0.06 |
| 6. Nacc tissue iron | -0.30* | -0.15* | -0.14* | -0.19* | 0.19* |  |  | 0.90 | 0.15 |
| 7. Caudate tissue iron | -0.04 | 0.10 | 0.04 | 0.07* | 0.34* | 0.06 |  | 0.91 | 0.08 |
| 8. Putamen tissue iron | -0.56* | -0.19* | -0.26* | -0.34* | 0.65* | 0.37* | 0.44* | 0.88 | 0.07 |

Note. Nacc = nucleus accumbens, *M* = mean, *SD* = standard deviation, \* = $p < 0.05$ . For tissue iron measures, negative correlations reflect positive relationships with other variables.

### MR data acquisition and preprocessing

#### Study A

For Study A, MRI data were acquired using a 7T Siemens scanner. High resolution structural images were acquired using a sequence that combines two fast gradient echo images with different inversion times (MP2RAGE; 1mm isotropic resolution, TI1/T2 (INV1/INV2), 800/2700 ms TR, 6000 ms; TE, 2.87ms; flip angle 1/2 (INV1/INV2), 4°/ 5°). Functional images were acquired via blood oxygen level dependent (BOLD) signal from an echo planar sequence (TR, 48 ms; TE, 23 ms; flip angle, 7°; voxel size at 2.0 mm isotropic resolution) with contiguous 2 mm – thick slices aligned to maximally cover cortex and basal ganglia. Participants completed an 8-minute fixation resting-state fMRI scan used for measurements of tissue iron, which given past studies (Larsen & Luna, 2015; Peterson et al., 2019), is sufficient for a reliable measure. Structural MRI data (T1w uniform images derived from the MP2RAGE acquisition) were preprocessed by skull extraction and linear (FLIRT) and non-linear (FNIRT) warping to MNI space. T2* data from resting-state fMRI underwent minimal preprocessing, including 4D slice-timing correction, head motion correction, skull stripping, co-registration to the structural image, and nonlinear warping to MNI space.

#### Study B

For Study B, MRI data were acquired using a 3T Siemens Biograph mMR PET/MRI scanner. High resolution structural images were acquired using T1-weighted magnetization-prepared rapid gradient-echo (MPRAGE) sequence (TR, 2300 ms; echo time (TE), 2.98 ms; flip angle, 9°; inversion time (T1) 900 ms; voxel size, 1.0 x 1.0 x 1.0 mm). Functional images were acquired via BOLD signal from an echoplanar sequence (TR, 1500 ms; TE, 30 ms; flip angle, 50°; voxel size, 2.3 x 2.3 x 2.3 mm in-plane resolution) with contiguous 2.3 mm – thick slices aligned to maximally cover cortex and basal ganglia. Participants completed two 8-minute (16 minutes total) fixation resting-state fMRI scans from which we derived T2* data. Structural MRI data were preprocessed by skull extraction and linear (FLIRT) and non-linear (FNIRT) warping to MNI space. T2* data from resting-state fMRI underwent minimal preprocessing, including 4D slice-timing correction, head motion correction, skull stripping, co-registration to the structural image, and nonlinear warping to MNI space.

#### nT2*w acquisition and preprocessing

As in previous work, tissue iron measures were obtained by normalizing and time-averaging T2* -weighted images (nT2*w) from echo planar imaging (EPI) scan sequences, which quantifies the relative T2* relaxation throughout the brain and is sensitive to magnetic field inhomogeneities caused by iron (Langkammer et al., 2010; Larsen & Luna, 2015; Peterson et al., 2019; Price et al., 2021), which is predominant in the basal ganglia.

Preprocessing steps for T2*-weighted images have been described elsewhere (Larsen & Luna, 2015; Parr et al., 2022; Peterson et al., 2019; Price et al., 2021; Vo et al., 2011). Briefly, each volume was z-score normalized using a coverage map from all non-zero valued inputs for each participant (Larsen & Luna, 2015; Peterson et al., 2019). The normalized signal was then aggregated voxel-wise across all volumes using the median, resulting in one normalized T2*-weighted (nT2*w) image per participant per available session. Volumes with frame-wise displacement (FD) > 0.3 mm were excluded (Siegel et al., 2014). This normalization process allows for the comparison of nT2*w values between-individuals (Larsen & Luna, 2015). Next, nT2*w values were extracted using for each basal ganglia region of interest (ROI) (globus pallidus, nucleus accumbens, putamen, and caudate nucleus) using the Harvard-Oxford atlas (Jenkinson et al., 2012). These values represent the mean across all voxels in each region, combining both hemispheres. Tissue iron (nT2*w values) was used in subsequent analyses as an indirect measure of basal ganglia dopamine neurobiology.

To control for differences in data acquisition effects between 3T and 7T scanners, nT2*w data were harmonized using neuroCombat (Fortin, 2024; Fortin et al., 2018). We note that nT2*w has a negative association with tissue iron; we therefore chose to reverse the direction of this measure in all axes in figures containing nT2*w to simplify interpretation. All statistical values reported in the text and tables were thus not modified and reflect numeric values.

### Two-stage sequential decision-making task

Participants completed a developmentally validated two-stage sequential decision-making task that has been used to differentiate model-based and model-free learning (Daw et al., 2011; Decker et al., 2016). Before completing the task, participants completed a tutorial in which they received instructions including the general cover story for the task, an explanation of probabilistic rewards and transitions, and completed practice trials.

The task has two stages with the goal of finding as much “space treasure” as possible (Figure 2A). First, participants can choose between two spaceships (first-stage choice). Each spaceship had a higher likelihood of traveling to one planet over another (70% versus 30%). For example, the blue spaceship had a 70% chance of reaching the red planet (common transition) and a 30% chance of reaching the purple planet (rare transition), whereas the green spaceship had a 70% probability of reaching the purple planet and a 30% probability of reaching the red planet. During the second stage, participants made a second choice between two alien stimuli (second-stage choice) that were rewarded with a “space treasure” (or nothing) according to a slowly drifting reward probability (ranging from 0.2 to 0.8). These changing reward probabilities encouraged participants to explore different second-stage options throughout the task to maximize their rewards. Participants had 3 seconds to make each choice, followed by a 1-second animation, 1-second of reward feedback, and a 1-second intertrial interval. The task comprised 200 trials divided into three blocks with optional breaks in between each block.

The structure of this task allows for the disassociation of model-based and model-free learning strategies. A model-based learner uses a cognitive model of transitions and outcomes to select actions, while a purely model-free learner repeats actions that were previously rewarded (Figure 2B). Thus, how the outcomes of previous trials influence subsequent first-stage choice depends on the learner’s strategy. For example, consider a situation where a participant chooses a blue spaceship, makes a rare transition to the purple planet, chooses an alien, and gets rewarded with “space treasure”. A model-free learner would likely repeat the blue spaceship choice and ignore the transition type (i.e., a main effect of reward). Conversely, a model-based learner would consider both the rewarded state and the state-transition structure of the task (i.e., that it was a rare transition), and thus would switch to the green spaceship on the next trial, regardless of the fact that they received a reward (i.e., a reward x transition interaction effect). Simply put, model-free learning strategies are biased towards recent rewards, and model-based learning is conceptualized as goal-directed control that integrates knowledge of task structure.

Building on model-based and model-free learning strategies, we derived a third measure that captures the frequency of each participant repeating the same choice of spaceship on subsequent trials regardless of transition-type (common versus rare transitions) and regardless of whether they earned a reward or not. This repetitive-response first-stage stay parameter can be thought to capture a participants’ tendency to respond habitually (i.e., perseveration), where behavior is outcome insensitive, independent from reward or transition-type, and are defined as persistence of behavior in the absence of the initial reward that established the stimulus response associations (Bouton, 2024).

#### Behavioral data preprocessing

Extending similar previous work, Multilevel logistic regression was used to estimate (1) model-based, (2) model-free, and (3) first-stage stay parameters for use in subsequent analyses. A detailed description of the multilevel logistic regression model for the two-stage sequential decision-making task has been described elsewhere (Daw et al., 2011; Decker et al., 2016; Otto et al., 2013, 2015). Briefly, the choice during the first stage (coded as stay = 1 or switch = 0 relative to the previous trial) was modeled by predictors of previous transition type (coded as common = 1 or rare = −1), previous reward outcome (coded as rewarded = 1 or not rewarded = −1), and their interactions as fixed effects. Random effects were estimated for the intercept, each predictor in the model (e.g., previous transition type and previous reward outcome), and their interaction for each available visit. The model-based parameter was calculated by adding the fixed effect of the interaction term and the random effect interaction term that was unique for each participant on each visit. The model-free parameter was calculated by adding the fixed effect of previous reward outcome term and the random effect of previous reward outcome that was unique for each participant on each visit. The first-stage stay parameter was calculated by adding the intercept term and the random intercept term that was unique for each participant on each visit. Trials in which participants failed to make a first- or second-stage choice were removed.

To validate our approach and replicate findings from prior developmental studies using the two-stage sequential decision-making task, we fitted an additional multilevel logistic regression model. In this model, age was included as a main effect, and all two- and three-way interactions were estimates, similar to Decker et al., (2016). Results from this analysis, which replicated past findings (Decker et al., 2016), are presented in Supplementary section S6 to allow for comparisons across developmental studies using this task. All multilevel logistic regression models were estimated using the lme4 package (Bates et al., 2015) in the R software environment (R Core Team, 2024).

We also fit participants’ choices to a set of three reinforcement learning (RL) models similar to the hybrid RL models described elsewhere (Daw et al., 2011; Wunderlich et al., 2012). These models consider the entire history of rewards and transition probabilities (as opposed to just previous trials in the multilevel logistic regression models) and assume that participant choices are driven by a weighted combination of model-based and model-free learning. The main goal of fitting RL models to the behavioral data was to compare the first-stage stay measure used in this study with a commonly estimated perseverance parameter from RL models (Katahira, 2018; Sugawara & Katahira, 2021). In this approach, the perseverance parameter reflects the tendency to repeat a first-stage choice regardless of the history of behavioral outcomes (i.e., outcome insensitive, indicating habitual behavior). Therefore, if the first-stage stay measure is indeed tracking habitual behavior, it should correlate with the perseverance parameter from the RL analysis. All reinforcement learning models were fit using the hierarchical Bayesian models of decision-making (hBayesDM) package in R (Ahn et al., 2017). Details of the model fitting procedure are presented in Supplementary section S7.

### Statistical Analyses

All statistical analyses were conducted using R Statistical Software (v4.4.1; R Core Team 2024). Zero-order correlations and descriptive statistics of all study variables are presented in Table 3.

To account for repeated measures, generalized additive mixed models (GAMM) were used, while also potentially capturing any non-linear effects in the developmental trajectories of nT2*w signal and the behavioral measures. All models included visit and sex assigned at birth as covariates. All GAMMs were fit using the *mgcv* package (Wood, 2017).

#### Characterizing developmental trajectories of nT2*w

We characterized developmental trajectories of tissue iron for each basal ganglia region (globus pallidus, nucleus accumbens, putamen, and caudate nucleus) similar to past studies from our group (Parr et al., 2021, 2022). Models assessing nT2*w developmental trajectories were specified as follows:

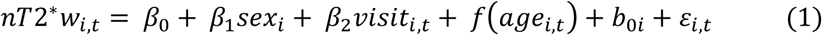

where *nT*2\**w_i,t_* represents the amount of iron for each ROI at visit *t* for person *i*; *β*_0_ is the intercept; *β*_1_ is the covariate for sex; *β*_2_ is the covariate for visit; *f*(*age_i,t_*) is a penalized smooth function for age with the maximum number of knots set to 3; *b_0i_* is the random intercept term for each participant, and 𝜀*_i,t_* is the residual error term for person *i* at visit *t*, which are assumed to follow a gaussian distribution. We applied the Bonferroni correction to account for multiple comparisons across the four ROIs, resulting in a significance threshold of α = 0.0125.

#### Characterizing developmental trajectories of behavioral performance

We characterized developmental trajectories of behavioral performance using GAMM models in a similar way as above. Models assessing developmental trajectories of behavioral performance during the two-stage sequential decision-making task were specified as follows:

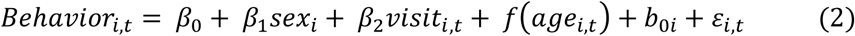

where *Behaviour_i,t_* represents measures derived from the multilevel logistic regression model (model-based, model-free, or first-stage stay) at visit *t* for person *i*. All other variables are specified and estimated identically as equation (1). We applied the Bonferroni correction to account for multiple comparisons across the three behavioral measures, resulting in a significance threshold of α = 0.0167.

To verify that the first-stage stay parameter tracked habitual responding, we examined the bivariate correlation between first-stage stays and the perseverance parameter created from the hBayesDM model (Ahn et al., 2017). Results from this analysis showed high correlation between the two parameters (*r* = 0.91) and are presented in Supplementary section S6.

#### Characterizing associations among tissue iron trajectories and behavioral performance

To examine associations among developmental trajectories of tissue iron across the four ROIs with behavioral performance, we compared a set of 4 models (Table 2), for each ROI separately, using the Akaike Information Criterion (AIC) and analysis of deviance tests to examine which model fit the data best (Wood et al., 2016). The four models increased in complexity, with the first model containing only a smooth term for age (identical to Equation 1), to a set of models where we added additional behavioral performance parameters. Our primary interest was to examine first-stage stays, because of its potential role in habits, however, we also considered the role of model-based and model-free control for a comprehensive evaluation of the task and for consistency with prior work (Decker et al., 2016). As such, we included the first-stage stay term first, then iteratively included model-based and model-free terms to see if they capture additional information about individual differences in tissue iron. This sequential model building approach allowed us to assess which behavioral parameters contribute the most to explaining variability in tissue iron trajectories. For each ROI, if the AIC and the analysis of deviance test indicated that a more complex model was a better fit, we retained that model and examined it further. All models that were used for comparisons were fit using maximum likelihood estimation. Model specifications for the analysis of deviance test and AIC comparisons are presented in Table 2.

**Table 2.**
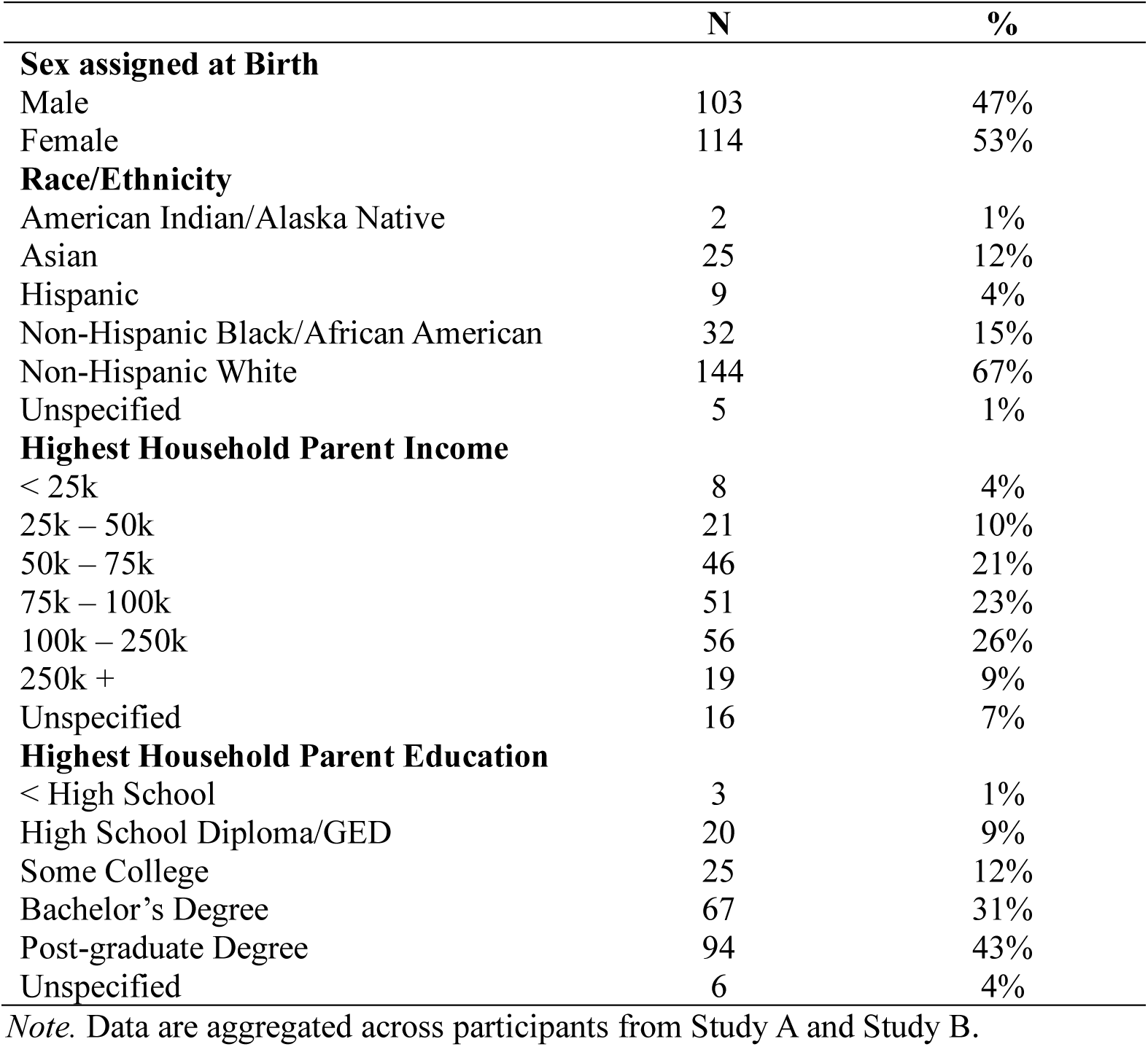
Baseline Demographic Information.

**Table 3.** Model specification of four different models to select from for associations among tissue iron and behavior.

| Model | Model specification |
| --- | --- |
| Model 1 | $nT2^*w_{i,t} = \beta_0 + \beta_1sex_i + \beta_2visit_{i,t} + f(age_{i,t}) + b_{0i} + \varepsilon_{i,t}$ |
| Model 2 | $nT2^*w_{i,t} = \beta_0 + \beta_1sex_i + \beta_2visit_{i,t} + f(age_{i,t}) + f(FSS_{i,t}) + b_{0i} + \varepsilon_{i,t}$ |
| Model 3 | $nT2^*w_{i,t} = \beta_0 + \beta_1sex_i + \beta_2visit_{i,t} + f(age_{i,t}) + f(FSS_{i,t}) + f(MB_{i,t}) + b_{0i} + \varepsilon_{i,t}$ |
| Model 4 | $nT2^*w_{i,t} = \beta_0 + \beta_1sex_i + \beta_2visit_{i,t} + f(age_{i,t}) + f(FSS_{i,t}) + f(MB_{i,t}) + f(MF_{i,t}) + b_{0i} + \varepsilon_{i,t}$ |

Note. FSS = first-stage stay; MB = model-based, MF = model-free.

#### Time-varying parameter models

To characterize how the relationship between tissue iron and habitual performance varied across development, we assessed whether the relationship between tissue iron and behavioral performance changed at different ages using the model that was identified via analysis of deviance test and AIC selection in the prior analysis. Time-varying parameter models allow for regression coefficients to be estimated as flexible, nonparametric smooth functions of age (Bringmann et al., 2017; Haslbeck et al., 2020; Hastie & Tibshirani, 1993; Sørensen et al., 2021; X. Tan et al., 2012). The time-varying parameter model was specified in this general form:

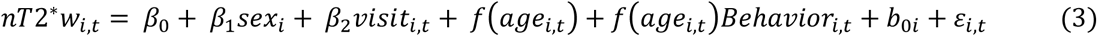

where *f*(*age_i,t_*)*Behaviour_i,t_* is a nonparametric term that allows the linear relationship between performance and iron to vary as a smooth function of age. To test for windows of significance across the age range, we calculated the first derivative of the time-varying smooth term from the GAMM via finite differences with 10,000 simulations using the *gratia* package in R. Significant differences were identified as age ranges where the 95% confidence interval of the derivative did not include 0.

## Data availability

Data will be made available on request.

## Code availability

Code for all figures and analyses are provided at https://github.com/LabNeuroCogDevel/daw_tissue_iron.

## Acknowledgments

This work was supported by grants 5R01MH080243 and 5R01MH067924 (A.C.P., F.C., B.L.) from the National Institutes of Mental Health, the Brain Behavior Research Foundation, and the Staunton Farm Foundation. We thank the University of Pittsburgh Clinical and Translational Science Institute (CTSI) for their support in recruiting participants, as well as their support by the National Institutes of Health through Grant Number UL1TR001857. Author D.J.P. was supported by the Developmental Alcohol Research Training Program (T32 AA007453) with funding from the National Institute on Alcohol Abuse and Alcoholism. Author V.J.S. was supported by NIMH T32MH016804.

## Contributions

D.J.P., A.C.P., B.T.C., F.C., and B.L. designed the study; D.J.P. and W.F. analyzed the data; D.J.P., V.S., and A.O. generated figures; D.J.P. wrote the original draft of the manuscript and all authors (D.J.P., A.C.P., V.S., A.O., W.F., B.T.C., F.C., and B.L.) reviewed and revised the final draft.

## Supplementary Materials

### >S.1. Supplementary results from section 3.2

Detailed summaries from GAMMs assessing developmental trajectories of tissue iron in the basal ganglia are presented in Supplementary Tables S1. All age smooths are significantly different than 0 at Bonferroni-adjusted α of 0.0125.

**Table S1.**
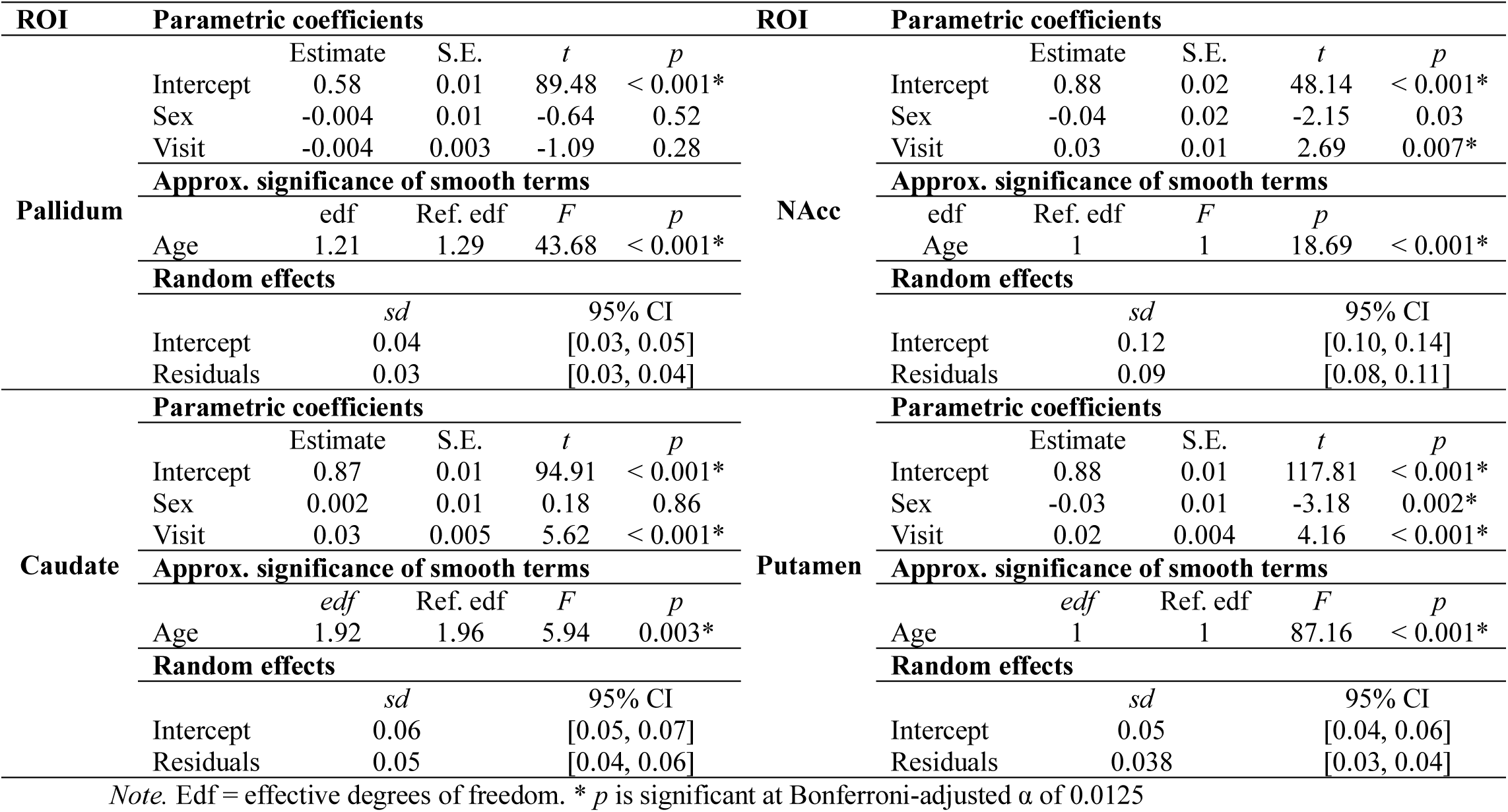
GAMM results characterizing developmental trajectories of nT2*w.

### >S.2. Supplementary results from section 3.3

Detailed summary tables from GAMMs assessing developmental trajectories of behavioral measures during the two-stage sequential decision-making task are presented in Supplementary Table S2. All age smooths are significantly different than 0.

**Table S2.**
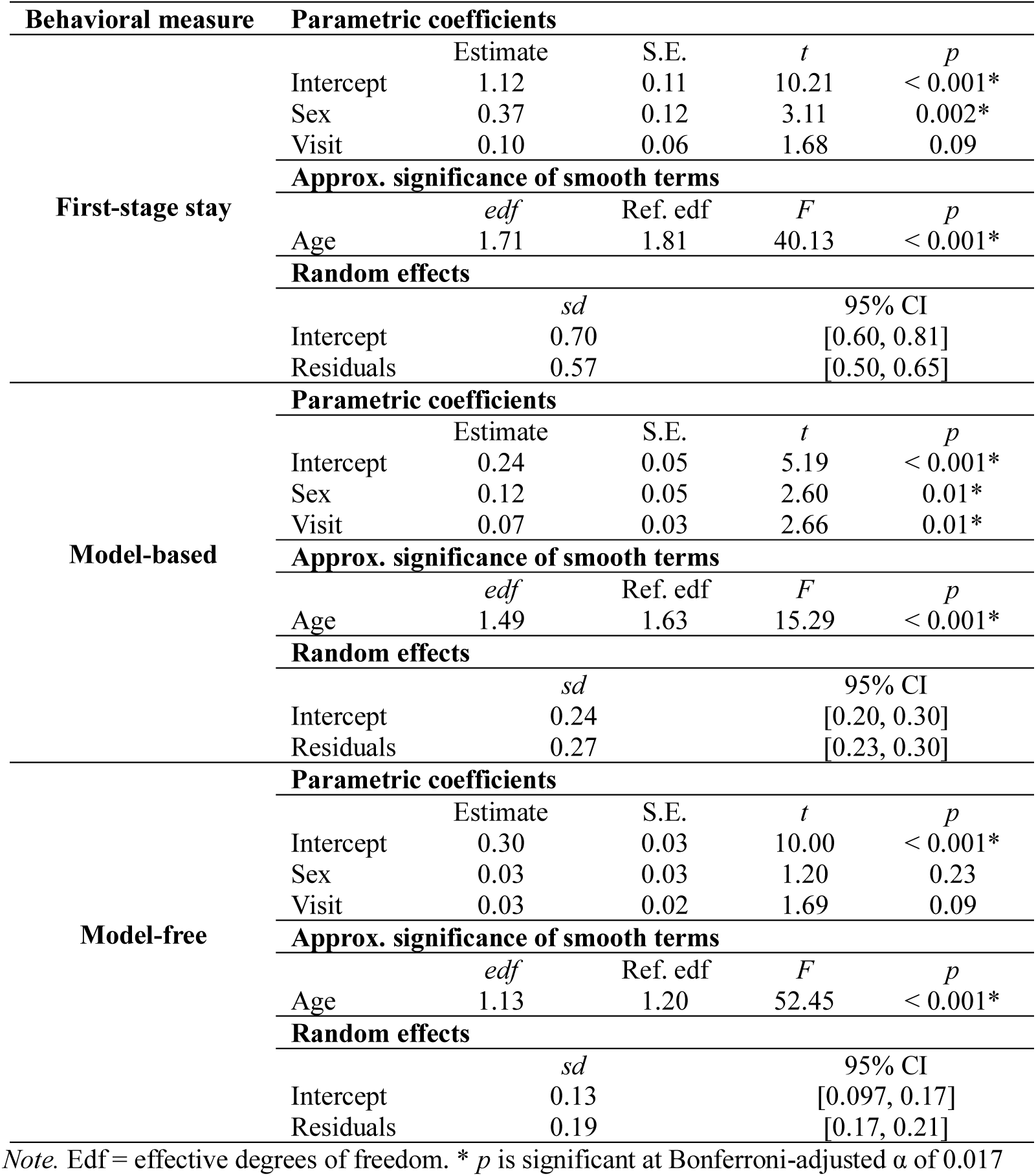
GAMM results characterizing developmental trajectories of behavioral measures.

### >S.3. Supplementary results from section 3.4

We compared a set of four models for each ROI using an analysis of deviance test and AIC comparisons (Table S3). For the caudate, NAcc, and globus pallidus, the analysis of deviance tests favored the simpler model with just a smooth term for age. For caudate and NAcc, the lowest AIC index was for model 1. For globus pallidus, the lowest AIC was for model 2. Thus, we did not retain any of these candidate models for further comparisons.

**Table S3.** AIC and analysis of deviance test for caudate, NAcc, and globus pallidus.

| ROI | Model | AIC | Resid. DF | Resid. Dev | DF | Deviance | *F* | *p* |
| --- | --- | --- | --- | --- | --- | --- | --- | --- |
| **Caudate** | **Model 1** | **-898.22** | 128.19 | 0.41 |  |  |  |  |
|  | Model 2 | -895.60 | 126.64 | 0.41 | 1.56 | -0.004 |  |  |
|  | Model 3 | -882.98 | 126.44 | 0.42 | 0.20 | -0.003 |  |  |
|  | Model 4 | -892.51 | 125.53 | 0.41 | 0.91 | 0.002 | 0.89 | 0.34 |
| **NAcc** | **Model 1** | **-495.71** | 122.55 | 1.44 |  |  |  |  |
|  | Model 2 | -495.37 | 120.57 | 1.43 | 1.98 | 0.006 | 0.33 | 0.72 |
|  | Model 3 | -493.82 | 119.97 | 1.44 | 0.60 | -0.001 |  |  |
|  | Model 4 | -492.15 | 119.40 | 1.44 | 0.59 | -0.003 |  |  |
| **Globus Pallidus** | Model 1 | -1126.39 | 122.23 | 0.19 |  |  |  |  |
|  | **Model 2** | **-1129.54** | 121.01 | 0.18 | 1.22 | 0.004 | 3.02 | 0.08 |
|  | Model 3 | -1128.94 | 120.19 | 0.18 | 0.82 | 0.0007 | 0.73 | 0.37 |
|  | Model 4 | -1128.41 | 117.24 | 0.18 | 2.95 | 0.0008 | 0.26 | 0.85 |
| **Putamen** | Model 1 | -1046.37 | 123.68 | 0.24 |  |  |  |  |
|  | Model 2 | -1048.81 | 122.88 | 0.24 | 0.80 | 0.003 | 3.16 | 0.09 |
|  | **Model 3** | **-1060.50** | 120.23 | 0.22 | 2.65 | 0.01 | 3.94 | 0.01* |
|  | Model 4 | -1058.09 | 119.85 | 0.23 | 0.38 | -0.001 |  |  |

*Note.* The lowest AIC fit index are bolded. AIC = Akaike information criterion. DF = Degrees of freedom for *F*-test, which reflect differences between degrees of freedom between models being compared. \* = $p < 0.05$ . For globus pallidus, the deviance test between model 1 and model 2 was not significant, indicating the model 1 was the better model.

Detailed summary tables from GAMM fits during model comparisons are presented in Supplementary Tables S4 – S7. For caudate (Table S5), NAcc (Table S6), and globus pallidus (Table S7), all age smooths were significant. However, no smooths from any of the behavioral measures were significant in the more complex models. For the putamen (Table S4), models 3 and 4 had significant smooths for age, first-stage stay, and model-based behavior.

**Table S4.**
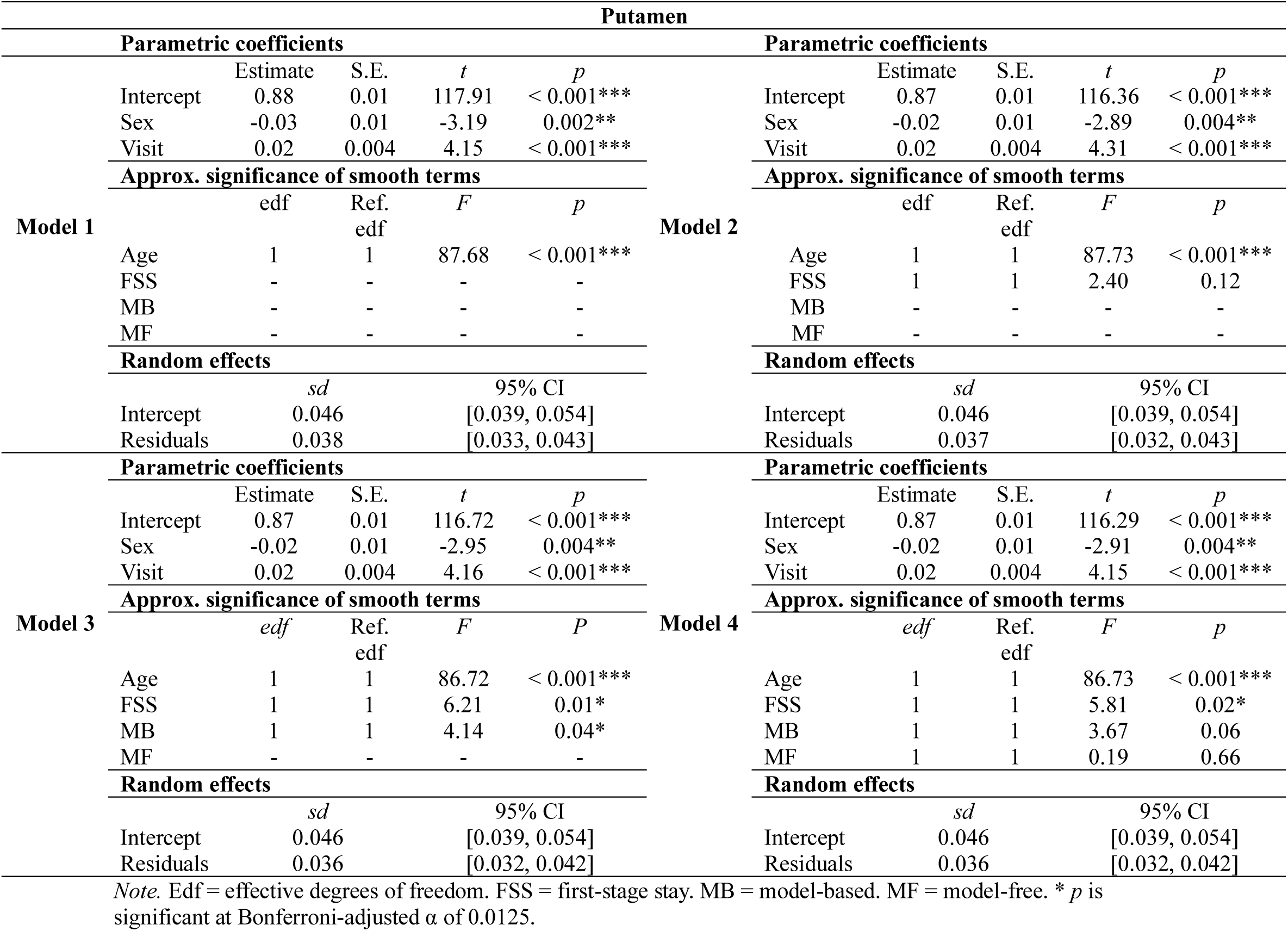
Full GAMM results of model comparisons for putamen.

**Table S5.**
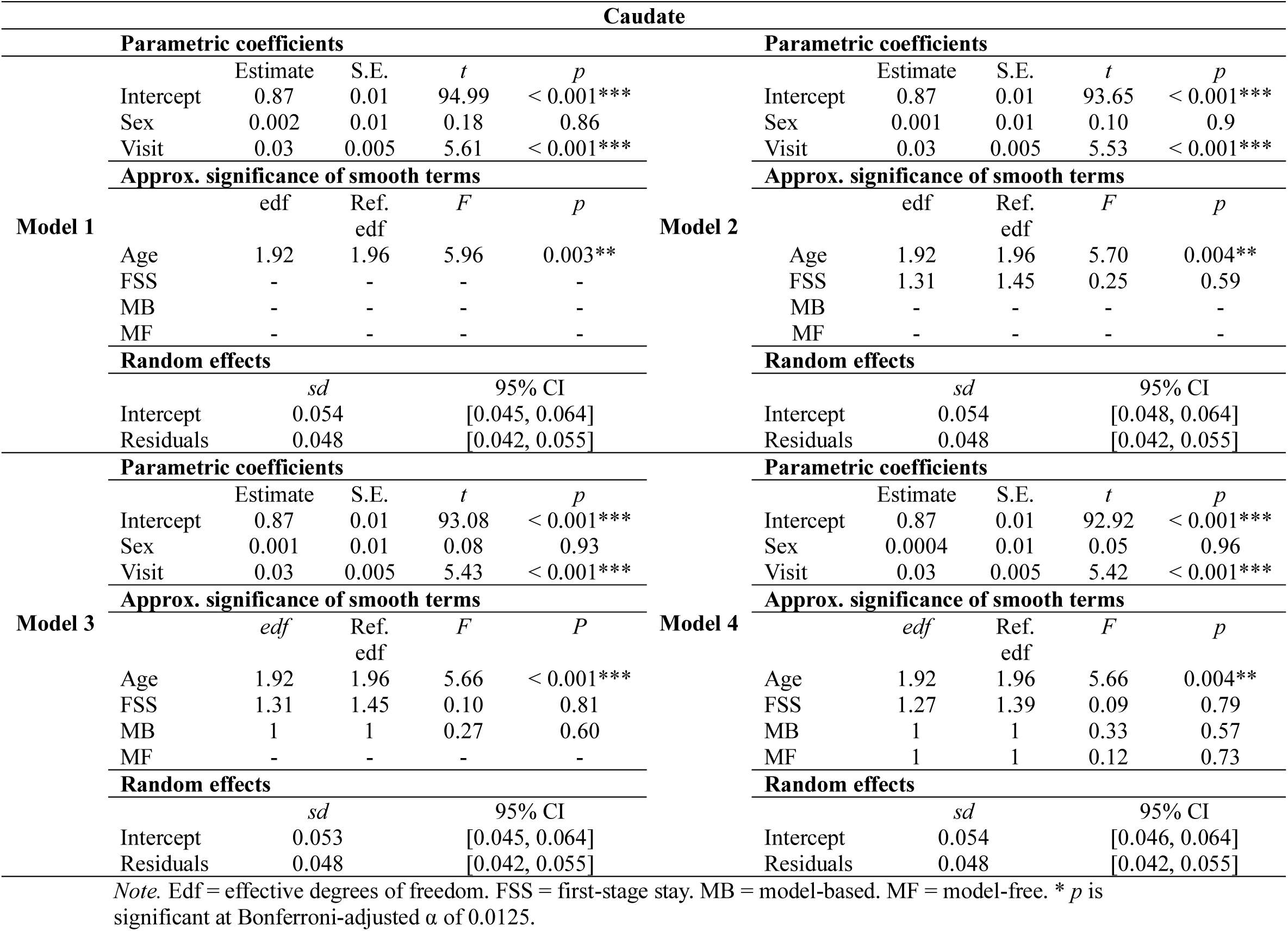
Full GAMM results of model comparisons from caudate.

**Table S6.**
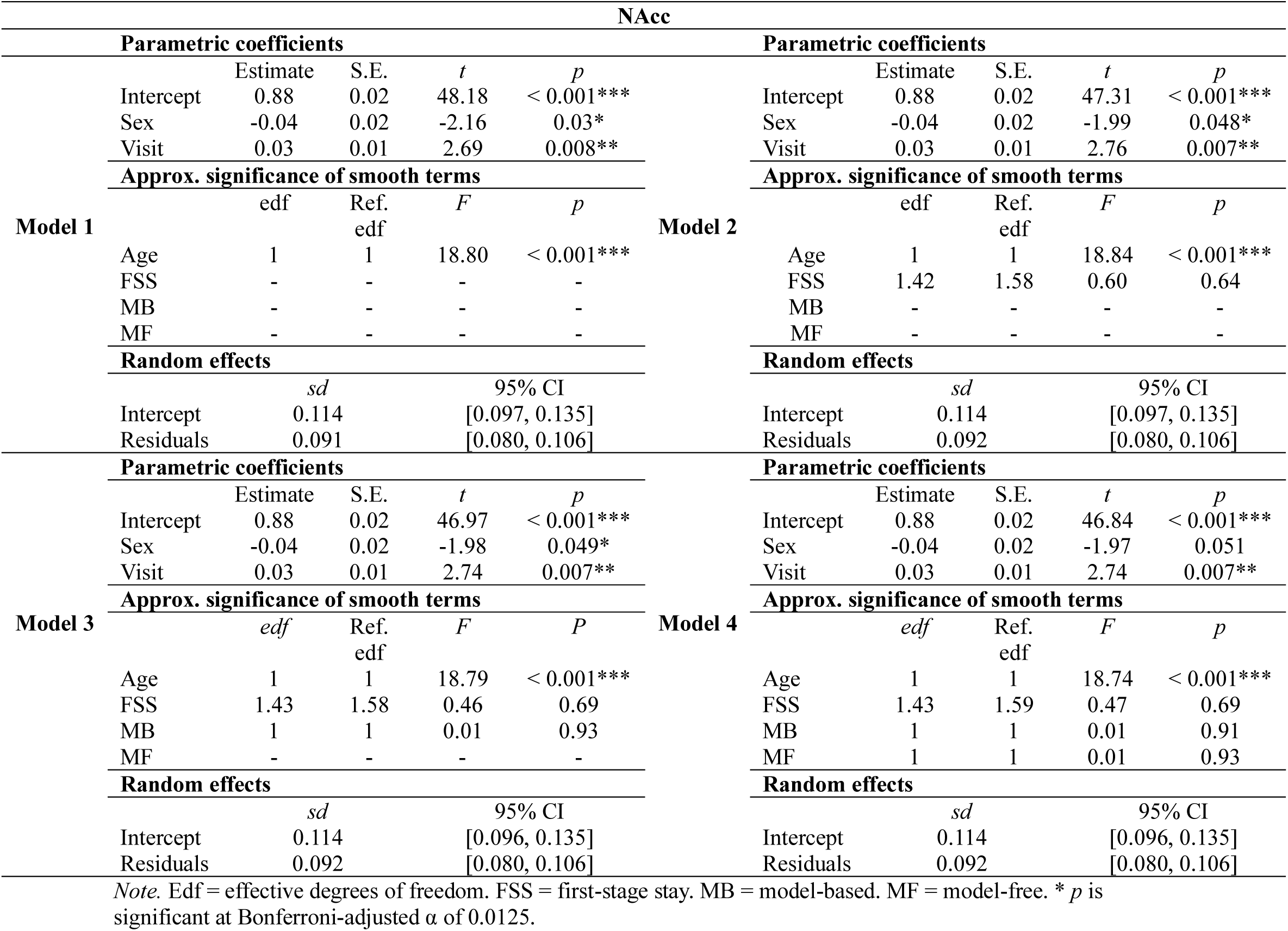
Full GAMM results of model comparisons from NAcc.

**Table S7.**
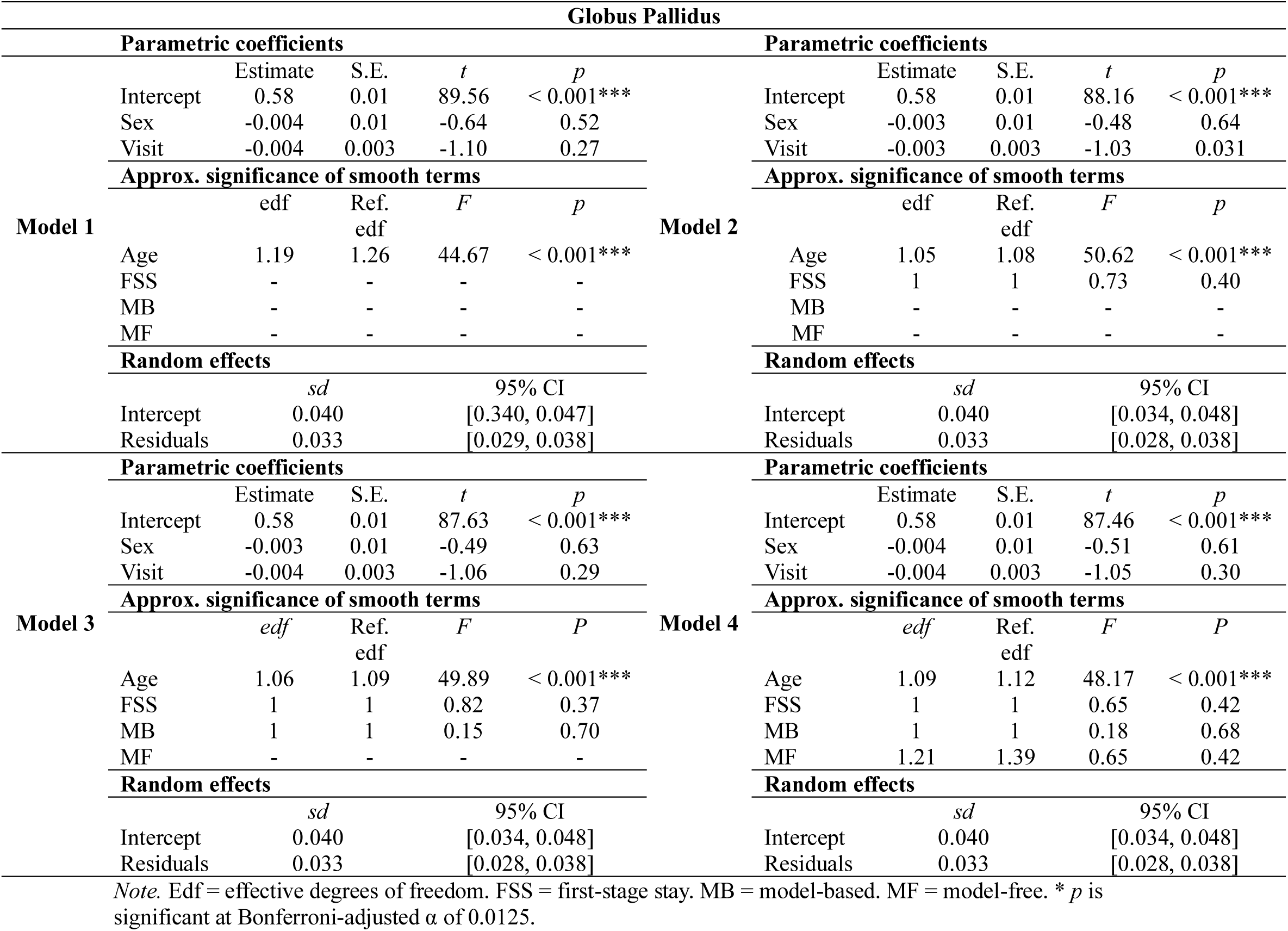
Full GAMM results of model comparisons from globus pallidus.

### S.4 Supplementary results from section 3.5

Detailed summary tables from GAMMs assessing whether developmental trajectories of tissue iron in the putamen varied with first-stage stay and model-based behavior (Table S8). Both time-varying parameters were significant, indicating that developmental trajectories of putamen tissue iron differed at varying levels of behavioral performance.

**Table S8.** Time-varying parameter modeling results.

| **Putamen** |  |  |  |  |
| --- | --- | --- | --- | --- |
| **Parametric coefficients** |  |  |  |  |
|  | **Estimate** | **S.E.** | ***t*** | ***p*** |
| Intercept | 0.87 | 0.01 | 117.63 | < 0.001*** |
| Sex | -0.02 | 0.01 | -3.10 | 0.002** |
| Visit | 0.02 | 0.004 | 4.23 | < 0.001*** |
| **Approx. significance of smooth terms** |  |  |  |  |
|  | **edf** | **Ref. df** | ***F*** | ***p*** |
| Age | 1 | 1 | 91.02 | < 0.001*** |
| Age:First stage stay | 2 | 2 | 4.34 | 0.014* |
| Age:Model based | 2 | 2 | 4.36 | 0.014** |
| **Random effects** |  |  |  |  |
|  | ***sd*** | **95% CI** |  |  |
| Intercept | 0.045 | [0.04, 0.05] |  |  |
| Residuals | 0.037 | [0.03, 0.04] |  |  |

Note. Edf = effective degrees of freedom. \* = $p < 0.05$ , \*\* = $p < 0.01$ , \*\*\* = $p < 0.001$ .

### S.5 Supplementary results from section 3.6

Detailed summary tables from GAMMs fit for the specificity analyses are presented in Supplementary Table S9.

**Table S9.**
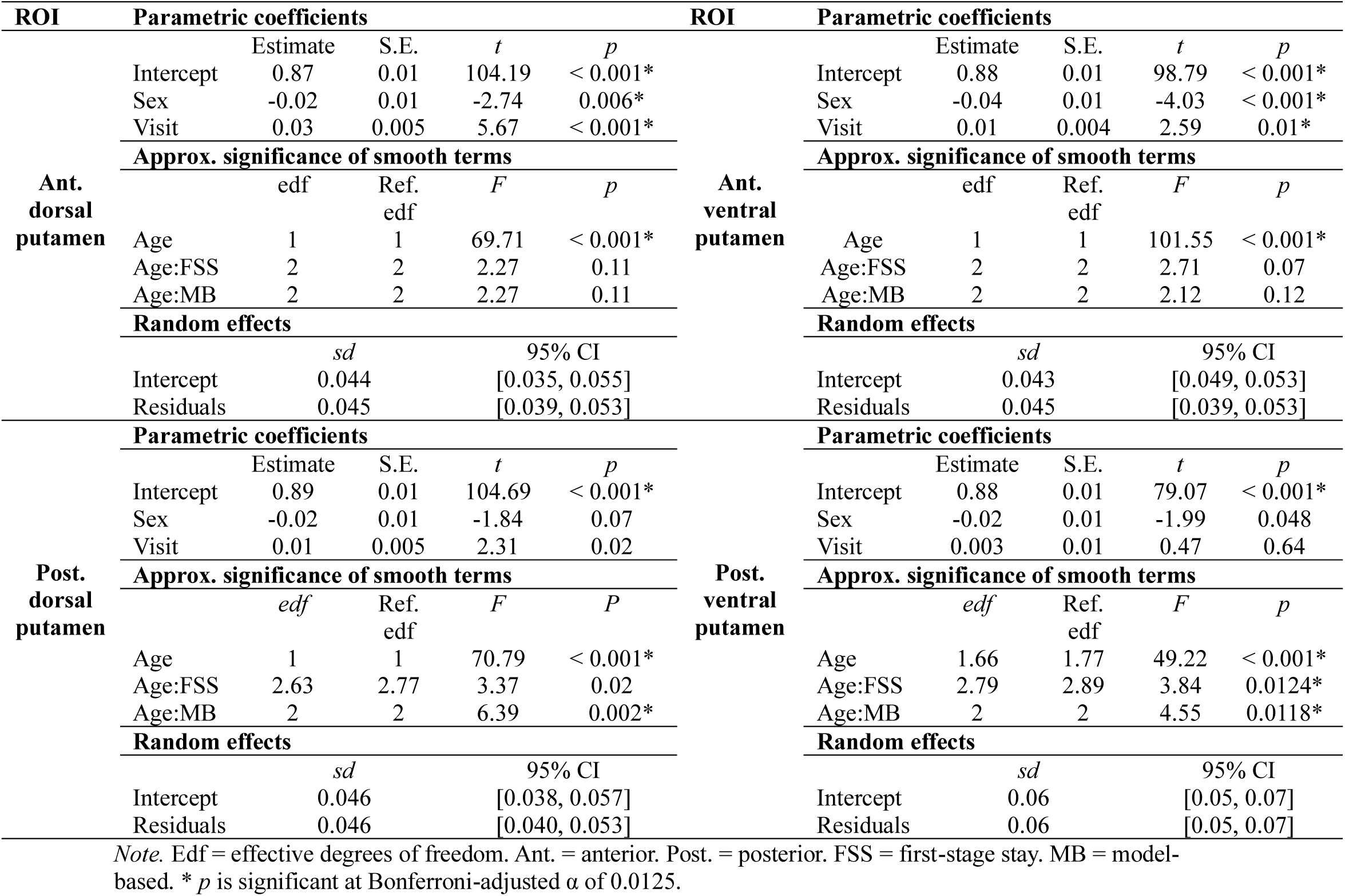
GAMM results characterizing developmental trajectories of putamen nT2*w according to CIC atlas.

**Figure S1.**
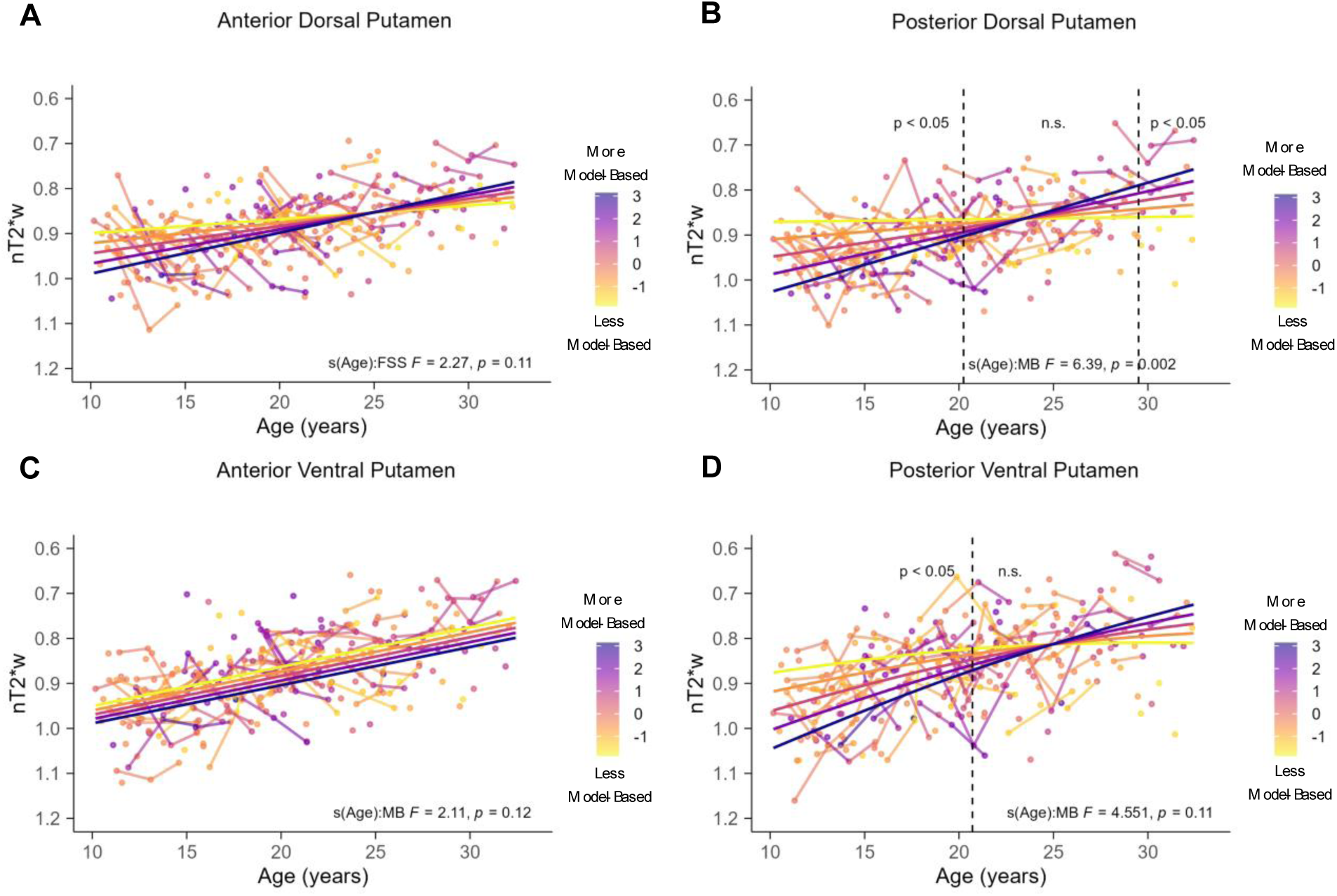
Time-varying parameter model results focusing on model-based behavior. In all panels, the y-axis is reversed such that lower values reflect more tissue iron. The color of the points represents model-based z-scores with darker colors indicating more model-based behavior. There were no significant associations between age and model-based behavior on anterior dorsal putamen tissue iron (A) or anterior ventral putamen tissue iron (C). (B) Model-based behavior moderated the association between age and posterior dorsal putamen tissue iron. The dashed vertical lines at 20.25 and 29.63 years old reflects the age boundaries where the 95% CI of the derivative does not include 0, indicating significance. (D) Model-based behavior moderated the association between age and posterior ventral putamen tissue iron. The dashed line at 20.71 years old reflects the age boundary where the 95% CI of the derivate does not include 0, indicating significance.

### S.6 Supplementary Behavioral Results

#### S.6.1 Multilevel logistic regression with continuous age

We tested for age related differences in the recruitment of each strategy using a continuous age term and its interaction terms, similar to the analyses conducted by Decker and colleagues (2016). The multilevel logistic regression model was specified as follows (using R syntax):

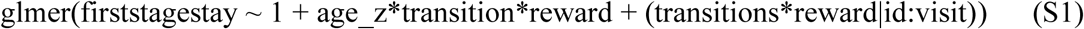

where the model was fit using the optimizer Bound Optimization by Quadratic Approximation (BOBYQA; Powell, 2009). Results from the multilevel logistic regression model are presented in Table S10. The effects of transition type and reward on the probability of a first-stage stay (FSS) faceted by age groups are presented in Supplementary Figure S2.

**Table S10.**
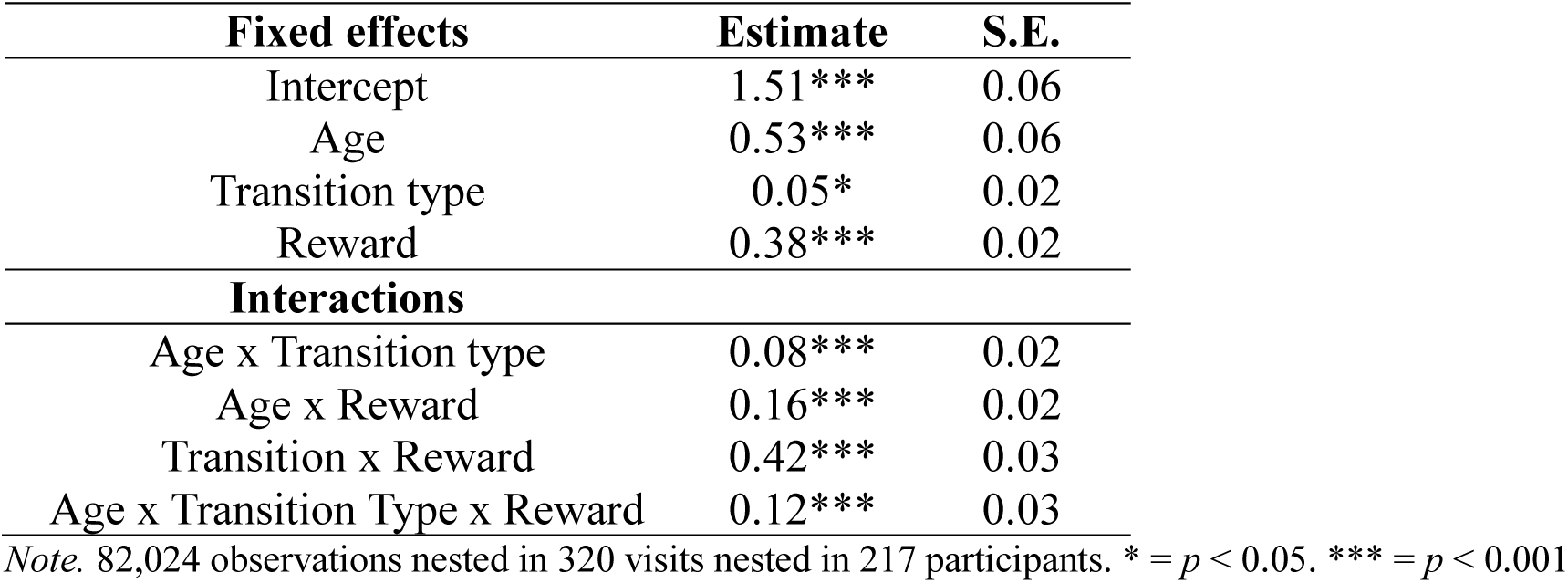
Multilevel logistic regression results.

| Fixed effects | Estimate | S.E. |
| --- | --- | --- |
| Intercept | 1.51*** | 0.06 |
| Age | 0.53*** | 0.06 |
| Transition type | 0.05* | 0.02 |
| Reward | 0.38*** | 0.02 |
| Interactions |  |  |
| Age x Transition type | 0.08*** | 0.02 |
| Age x Reward | 0.16*** | 0.02 |
| Transition x Reward | 0.42*** | 0.03 |
| Age x Transition Type x Reward | 0.12*** | 0.03 |

Note. 82,024 observations nested in 320 visits nested in 217 participants. \* = $p < 0.05$ . \*\*\* = $p < 0.001$

**Figure S2.**
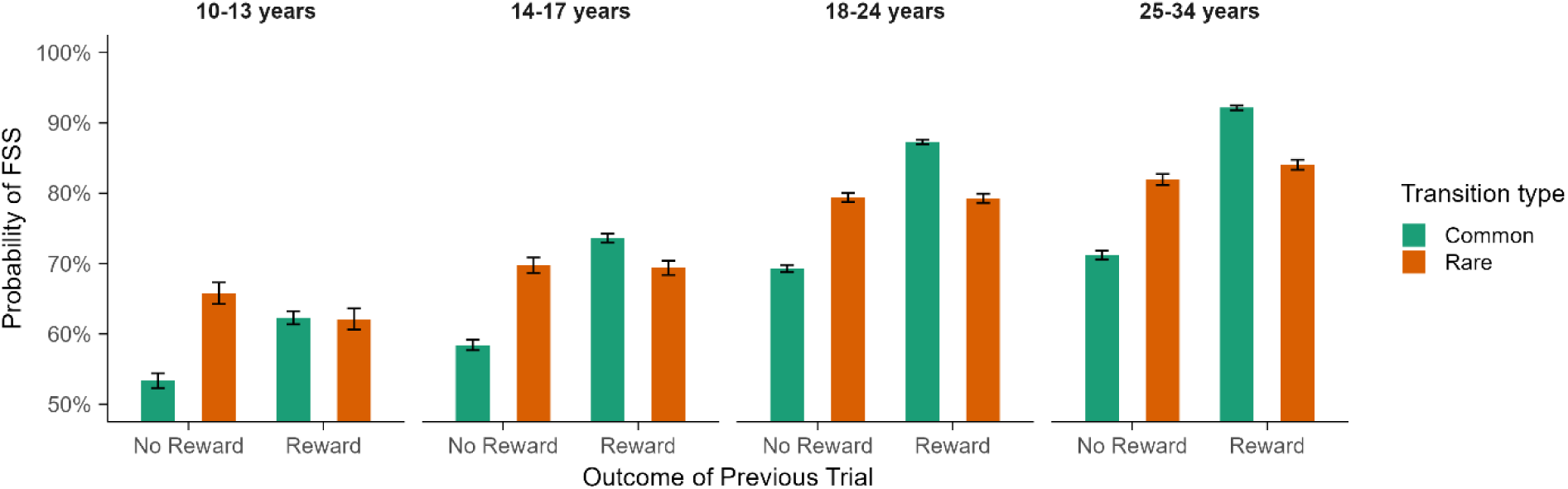
Probability of first-stage stays, averaged across participants and binned by representative age groups. Error bars represent +/- 1 SEM. Across all age groups, there is evidence of participants using both model-based and model-free strategies.

The main effects of age, transition type, and reward were significant, indicating that as age increased, the probability of a FSS increased (Figure S3 Panel A.), the probability of a FSS was greater after common transitions (Figure S3 Panel B.), and the probability of a FSS was greater after earning a reward (Figure S3 Panel C.). All two-way interactions were significant. We found that transition type moderated the association between age and FSS (Figure S3 Panel D), such that as age increased common transitions were associated with more FSS. We found that reward moderated the association between age and FSS (Figure S3 Panel E), such that in young participants the difference between receiving a reward and not were less associated with FSS, in older participants that difference was more pronounced, in that older adults had more FSS after receiving a reward. We also found that reward moderated the association between transition type and FSS (Figure S3, Panel F), such that the highest probability of a FSS occurred after being rewarded after a common transition. We also observed a significant three-way interaction (Figure S4). After rare transitions, there were no differences between earning and not earning a reward across the entire range of age, whereas after common transitions, the difference between receiving a reward and not were less associated with FSS, compared to older participants where receiving a reward was associated with more FSS. Taken together, these findings suggest that model free (main effect of reward) and model based (transition type x reward interaction) behaviors are evident in the full sample, albeit to differing degrees. Adults appear to be using all strategies at a higher level than adolescents.

**Figure S3.**
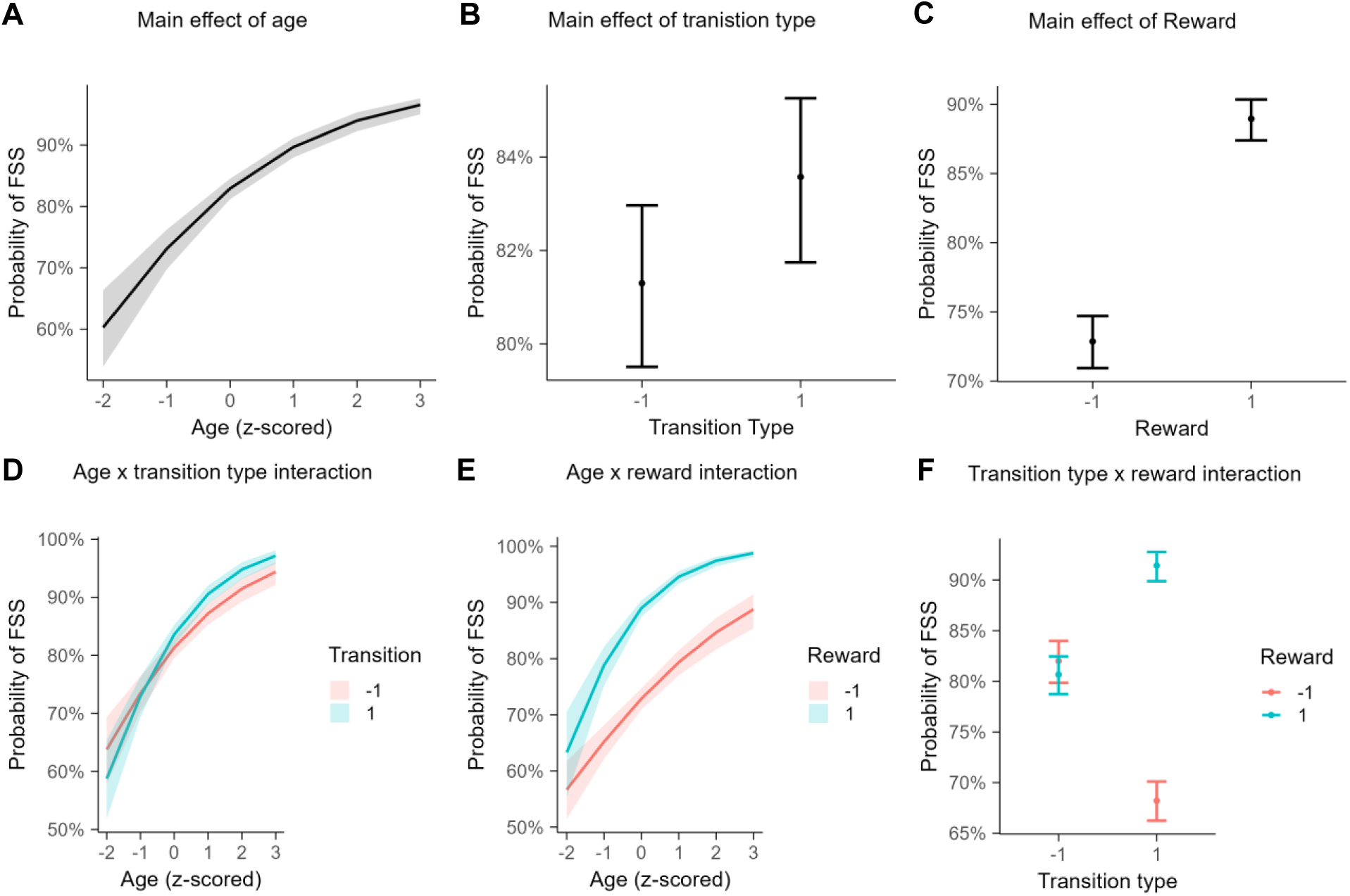
Marginal effects plots from the multilevel logistic regression model with continuous age. (A.) We observed a main effect of age, such that older participants had a higher probability of a FSS compared to younger participants. (B.) We observed a main effect of transition type, such that common transitions were associated with a higher probability of a FSS compared to rare transitions. (C.) We observed a main effect of reward (i.e., model-free behavior), such that receiving a reward was associated with a higher probability of a FSS compared to not being rewarded. (D.) We observed an age by transition type interaction, such that the probability of a FSS was greater after a common transition compared to a rare transition for adults but not for adolescents. (E.) We observed an age by reward interaction, such that the difference in the probability of a FSS after receiving versus not receiving a reward was the most pronounced for older participants compared to younger participants. (F.) We observed a transition type by reward interaction (i.e., model-based behavior), such that after rare transitions, there was minimal differences in the probability of a FSS after receiving verses not receiving a reward. For common transitions, the probability of a FSS was the highest after receiving a reward.

**Figure S4.**
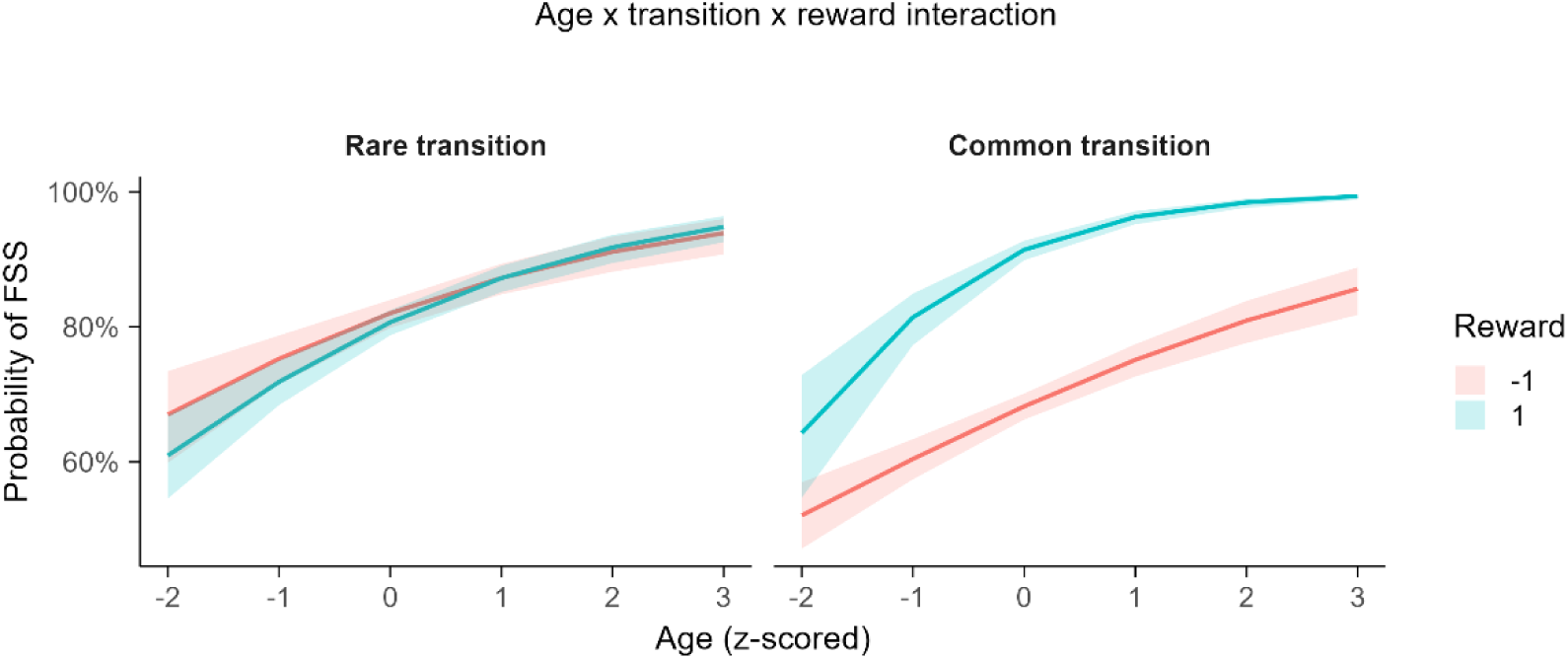
Marginal effects plot of the three-way interaction from the multilevel logistic regression model with continuous age. The probability of a FSS after a rare transition increased with age, but was not different depending on whether participants earned a reward or not. The probability of a FSS after a common transition also increased with age. However, the age increase in FSS was pronounced after being rewarded compared to not being rewarded.

#### S.6.2 Relation between knowledge of task structure and learning

Similar to other developmental studies using the sequential decision-making task (Decker et al., 2016; Nussenbaum et al., 2020; Potter et al., 2017), we assessed whether participant responses reflected knowledge of task structure, and whether knowledge of task structure was related to model-based control.

To assess participants’ knowledge of task structure, we examined whether participants’ response times when making second-stage choices were affected by the type of transition they experienced. If participants were unaware of the task structure, we would expect no difference in RTs following common versus rare transitions. Conversely, if they understood the transition structure, we would anticipate slower responses after rare transitions. A linear mixed-effect model was used to examine these associations (see Table S11 and Figure S5A). Results suggest that participants were slower after rare transitions compared to common ones (β = −41.27, SE = 1.47, *p* < 0.001), that older participants made faster choices (β = −31.38, SE = 8.49, *p* < 0.001) and that the effect of transition type on reaction times increased with increasing age (β = −20.98, SE = 1.47, *p* < 0.001).

To assess whether knowledge of task structure was related to model-based control, we fit an additional linear multilevel model where we computed a reaction time difference score for the second stage choice by subtracting each participant’s mean reaction time after common transitions from each participant’s mean reaction time after rare transitions. Then, we regressed the model-based coefficient on age, the mean difference in reaction times during the second stage choice, and their interaction. We found the reaction time difference scores were associated with model-based control (β = 0.15, SE = 0.02, *p* < 0.001), indicating that slower reaction times after rare transitions were associated with more model-based control. We did not observe a significant reaction time difference by age interaction when we treated age as a continuous variable. Full results are presented in Table S12 and Figure S5B. Taken together, these results suggest that participants did understand the transition structure of the task. However, early adolescents do not appear to be integrating this information to the extent that mid/late adolescents and adults do. Our behavioral results are generally consistent with other research groups who have administered this task to developmental samples.

**Figure S5.**
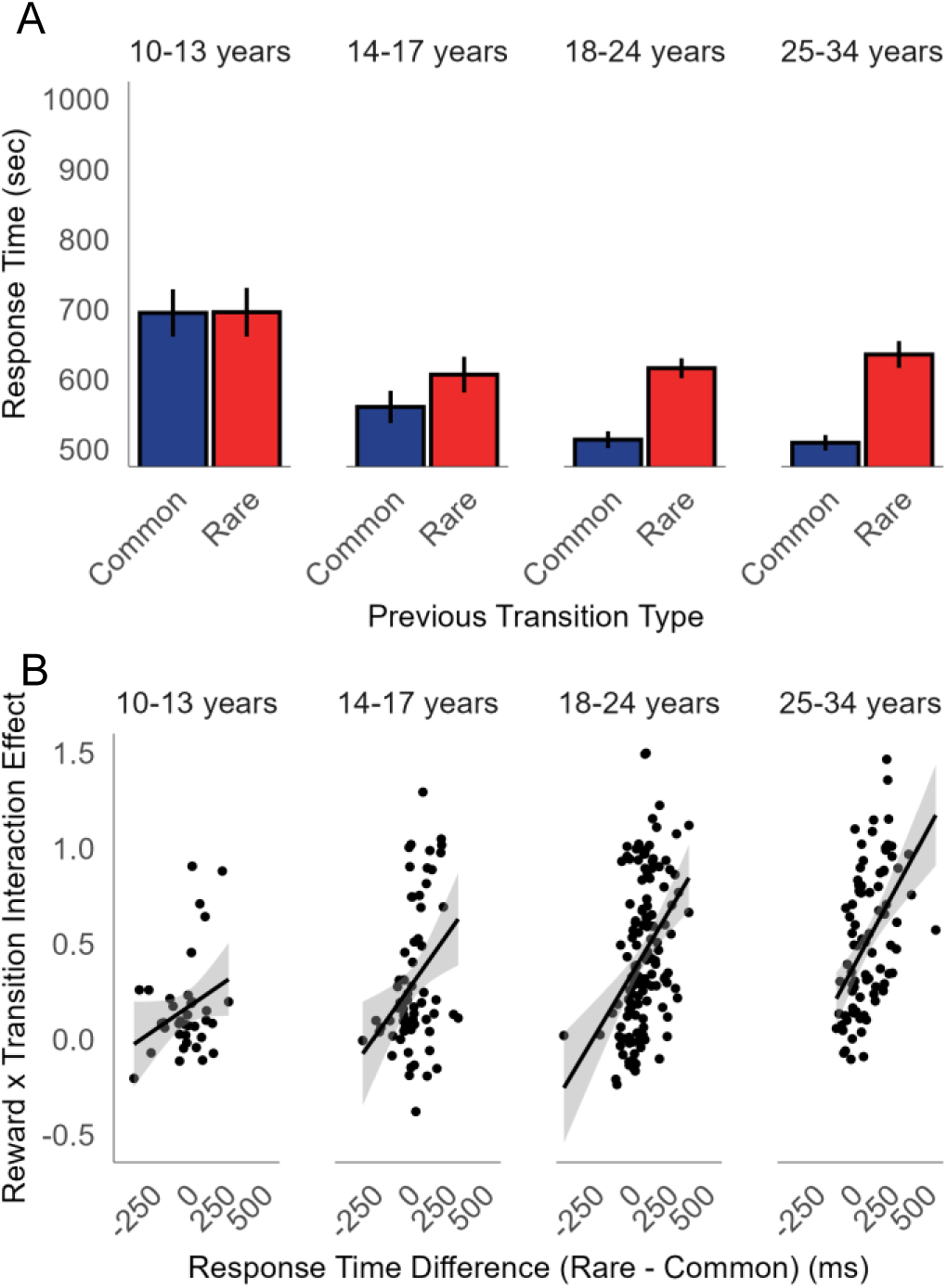
Second-stage response time results. (A) Bar graphs depicting response times for choices during the second-stage as a function of the previous transition type for each age group. Error bars reflect ± 1 *SEM*. On average, participants were slower during their second-stage choices following rare transitions compared to common transitions. This difference in response times between transition types increased across age. (B) Scatter plots depicting the relationship between the difference in response times between rare and common transitions and the reward x transition interaction effect (i.e., model-based term) for each age group. On average, participants with larger response times after rare transitions compared to common transitions had a larger reward x transition interaction effect. This effect increased across age. These results suggest that knowledge of task structure was related to participants’ use of model-based control.

**Table S11.** Linear multilevel modeling coefficients indicating the effect of age and transition type on second-stage choice reaction times.

| **Fixed Effects** | **Estimate** | **SE** | ***p*** |
| --- | --- | --- | --- |
| Intercept | 583.55 | 8.50 | < 0.001 |
| Age | -31.38 | 8.49 | < 0.001 |
| Transition type | -41.27 | 1.47 | < 0.001 |
| Age x Transition type | -20.98 | 1.47 | < 0.001 |
| **Random Effects** | ***sd*** | **95% CI** |  |
| Intercept | 149.70 | [138.08, 161.89] |  |
| Residuals | 342.4 | [340.48, 344.23] |  |

**Table S12.** Linear multilevel modeling coefficients indicating the effect of age and RT differences on model-based control.

| **Fixed Effects** | **Estimate** | **SE** | ***p*** |
| --- | --- | --- | --- |
| Intercept | 0.38 | 0.02 | < 0.001 |
| Age | 0.07 | 0.02 | 0.0014 |
| RT difference | 0.15 | 0.02 | < 0.001 |
| Age x RT difference | 0.01 | 0.02 | 0.48 |
| **Random Effects** | ***sd*** | **95% CI** |  |
| Intercept | 0.21 | [0.16, 0.26] |  |
| Residuals | 0.26 | [0.23, 0.30] |  |

*Note.* RT = Reaction Time

### S.7 Reinforcement learning model details

The primary purpose of fitting RL models to the behavioral data was to compare the first-stage stay measure used in the current study to a commonly estimated perseverance parameter that is estimated in some RL models. Briefly, the perseverance parameter tracks the tendency to repeat a first-stage choice regardless of outcome (i.e., outcome insensitive, which is indicative of habits). Thus, if the first-stage stay measure is tracking habitual behavior, then it should correlate with the perseverance parameter. The following section describes the set of RL models used for this comparison in more detail.

#### S.7.1 hBayesDM

Hierarchical Bayesian analyses (HBA) were conducted using the hierarchical Bayesian Decision-Making (hBayesDM) package for R (Ahn et al., 2017) that uses rstan (version 2.32.6) for implementation. In Bayesian statistics, we start with prior beliefs (known as prior distributions) about the model parameters. These priors are updates into posterior distributions based on the data (i.e., trial-by-trial choice and outcomes during the two-stage task) by applying Bayes rule (Kruschke, 2014). In a HBA, group or hyperparameters are introduced in addition to the individual-level parameters which leads to shrinkage effects (Gelman et al., 1995). Shrinkage effects in a HBA occur when each individual’s estimates influence the group’s overall estimates, which then feedback to refine the estimates for each individual. A result of shrinkage effects is that individual estimates are often more stable and reliable with this approach, because similarities among participants are informed by the group parameters. Posterior inference for hBayesDM models use a Markov chain Monte Carlo (MCMC) sampling scheme called Hamiltonian Monte Carlo (HMC; (Carpenter et al., 2017). A full description of hBayesDM can be found in Ahn et al. (2017).

#### S.7.2 Two-stage sequential decision-making parameter descriptions

The hBayesDM packages come with three pre-programmed RL models based on RL models described in Daw et al. (2011) and Wunderlich et al., (2012). Each model differs in the number of free parameters that are estimated (either 4 parameters, 6 parameters, or 7 parameters). All RL models contain the following free parameters (α, β, ᴨ, w):

The alpha (α) parameter (learning rate) determines how quickly an agent updates their expectation based on the prediction error (e.g., the difference between expected and actual outcomes) at both the first and second stages. A higher learning rate means the agent rapidly adapts to new information, while a lower learning rate indicates a more conservative updating of beliefs. The beta (β) parameter (inverse temperature) controls the balance between exploration and exploitation in decision-making. A higher beta value means the agent is more deterministic in their choices, favoring the option with the highest expected value, implying more exploitation. A lower beta suggests greater randomness in choice, implying more exploration. The pi (ᴨ) parameter (perseverance) reflects the tendency of the agent to repeat a previous choice regardless outcome. It captures how “sticky” the agent’s bias is towards a previously chosen action. Higher values indicate a stronger tendency to persevere with the same choice. The W (w) parameter (model-based weight) represents the weight given to model-based versus model-free behavior.

Model-based behavior involves planning and using a cognitive map of the task structure to make decisions, while model-free behavior relies on cached values from past experiences. A higher w indicates greater reliance on model-based strategies, whereas a lower w suggests more dependence on model-free learning.

The only difference between the 4-parameter model and the 6-parameter model is that α and β are estimated for each stage of the task (e.g., α stage 1/ α stage 2, β stage 1/ β stage 2). For the 7-parameter model, a lambda (λ) free parameter is estimated. Lambda is used in temporal difference learning to determine how much of the prediction error is assigned to past states and actions. It influences the extent to which past experiences (not just the most recent one) are updated based on new information. A higher lambda means that more distant past experiences are considered when updating value.

#### S.7.3 Computational model description

For the full description of the computational model, see Daw et al., (2011). hBayesDM code for the two-stage sequential decision-making task is from the publicly available GitHub repository (https://github.com/ccs-lab/hBayesDM). Briefly, the task has three states (first stage: 𝑠_*A*_; second stage: 𝑠_*B*_ and 𝑠_*C*_) and two actions (*a_A_* and *a_B_*). The temporal difference learning algorithm learns a state-action value function *Q_𝑠,a_* by mapping each state and action to its expected future value by an update rule for stage *i* for each trial *t* according to (notation is from Daw et al. (2011):

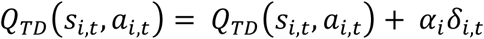

where

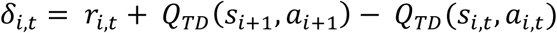

and *α_i_* is the learning rate parameter. *δ_i,t_* reflects the reward prediction error which is driven by the second-stage value only (because *r_i,t_* = 0). The λ parameter effects first stage action by the second stage reward prediction error according to:

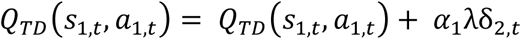

The w parameter is introduced to connect values to choices by a weighted sum of model-based and model-free values according to:

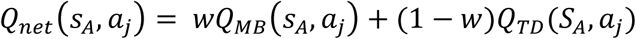

Finally, the probability of a choice is calculated via a softmax function as:

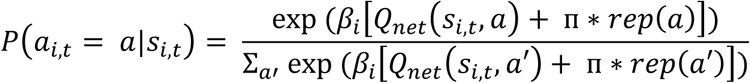

where *β_i_* is the inverse temperature parameter that control how deterministic or stochastic the choices are and ᴨ is the perseverance parameter that controls for how sticky first stage choices are.

#### S.7.4 Model selection

We fit the three described models on the behavioral data, each with 4 chains, 1000 burn-in samples, and 3000 samples. Model selection was conducted using Leave-One-Out Information Criterion (LOOIC) and Widely Applicable Information Criterion (WAIC; see Vehtari et al., 2017 for details). Lower LOOIC and WAIC indicate better model fit. The 7-paramter model was found to be the winning model and was used for further analyses.

**Table S11.** Global fit statistics used for model selection.

| Model | LOOIC | WAIC |
| --- | --- | --- |
| 4-parameter model | 165142.20 | 164181.40 |
| 6-parameter model | 163580.30 | 162760.00 |
| 7-parameter model | **163545.50** | **162754.00** |

Note. Lowest LOOIC/WAIC are made bold.

#### S.7.5 Assessing model convergence

We examined *R*^ and visually diagnosed MCMC performance to assess model convergence. *R*^ values for the hyper parameters were all close to 1, indicating convergence (*R*^ 1 – 1.01). For individual parameter estimates (including the posterior prediction estimates), only five individual parameters had an *R*^ value that exceeded 1.04. For the visual diagnosis, we checked whether MCMC samples were well mixed and converged to stationary distributions.

#### S.7.6 Bivariate association between first-stage stay and perseverance parameters

We assessed the bivariate association between first-stage stay behavior derived from the multilevel logistic regression and perseverance parameter from the best fitting RL model. If first-stage stay behavior is indicative of habitual behavior, then it should be correlate with the perseverance parameter. Additionally, the perseverance parameter should have a similar developmental trajectory compared to the first-stage stay measure. Results from the supplemental analysis found that the correlation between first-stage stay and perseverance was high (Figure S6 A; *r* = 0.91). Both measures also had similar developmental trajectory patterns, with increases across the age range (Figure S6 B). Thus, it is reasonable to suggest that first-stage stay is tracking habitual responding during the two-stage sequential decision-making task.

**Figure S6.**
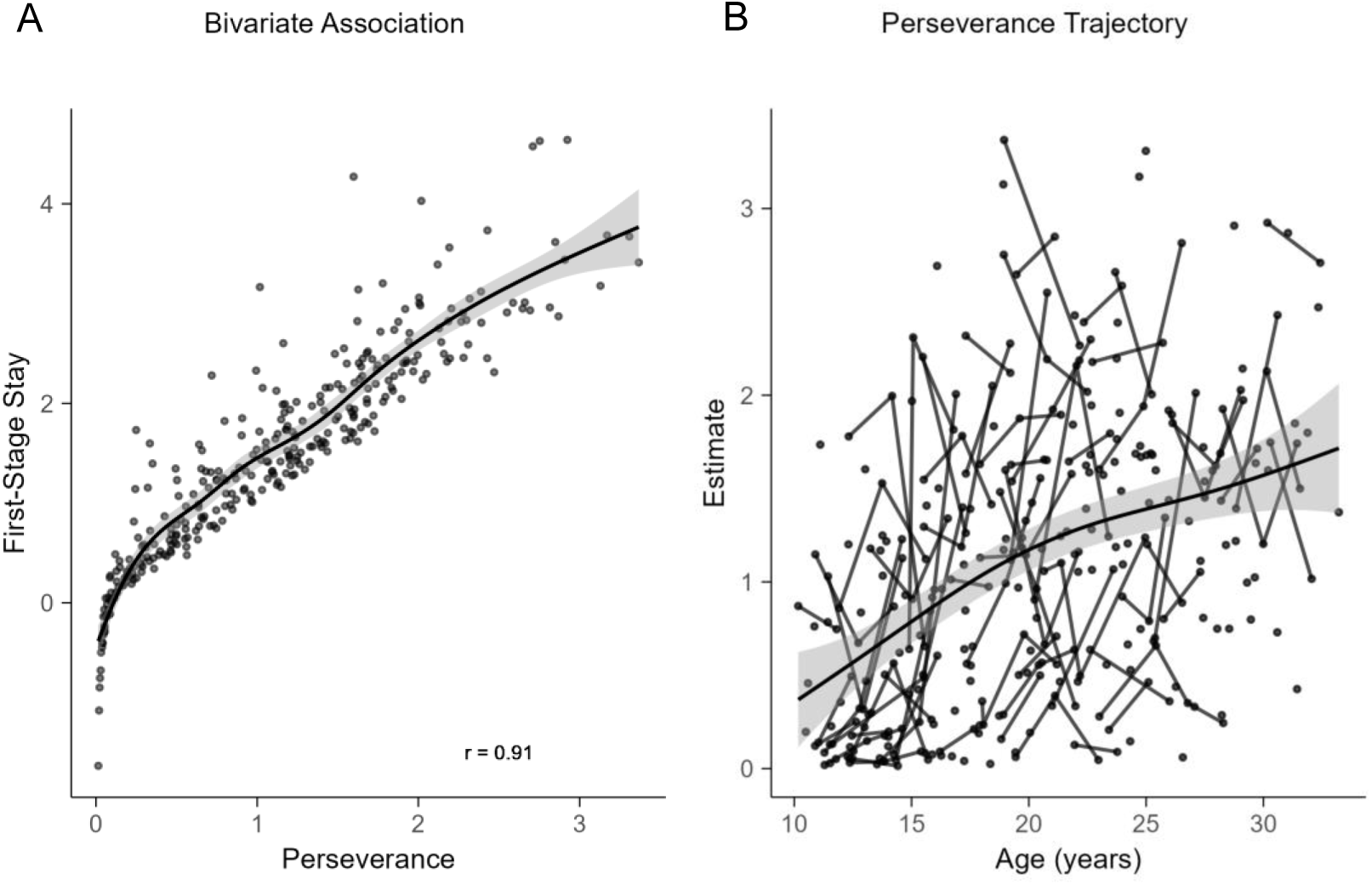
Plots comparing the first-stage stay measure to the perseverance parameter. (A) The bivariate correlation between perseverance and first-stage stay is large (*r* = 0.91) indicating that they measure a similar latent construct (i.e., habitual responding). (B & C) We observed similar age trajectories for the first-stage stay measure and the perseverance parameter.

